# Inter-Spheroid Proximity and Matrix Remodeling Determine CAF-Mediated Cancer Cell Invasion

**DOI:** 10.1101/2025.04.24.648914

**Authors:** Pranav Mehta, Ankur Deep Bordoloi, Cor Ravensbergen, Ma. Kristen H. David, Wilma Mesker, Gerrit Jan Liefers, Peter ten Dijke, Pouyan E. Boukany

## Abstract

Breast cancer is the most commonly diagnosed malignancy worldwide, with molecular subtypes following distinct clinical trajectories. While Luminal A breast cancers are typically indolent, a subset enriched in α-smooth muscle actin (α-SMA)-positive cancer-associated fibroblasts (CAFs) exhibits aggressive behavior, facilitating tumor invasion. However, the biophysical mechanisms by which CAFs drive invasion and extracellular matrix (ECM) remodeling remain unclear. Furthermore, the temporal and spatial dynamics of CAF interactions with the collagen matrix and cancer cell spheroids remain unknown, raising the question of whether these processes follow a deterministic sequence or occur stochastically. To address this, we conduct histological analysis of Luminal A tumors, revealing variation in CAF, cancer cell, and ECM organization at tumor boundaries. To assess the impact of CAF on cancer cell invasion, we use a 3D *in-vitro* model co-embedding 19TT breast CAF and MCF7 luminal breast cancer spheroids within a three-dimensional (3D) collagen-I hydrogel and perform time-lapse imaging. We demonstrate that inter-spheroid distance critically determines 19TT CAF-induced MCF7 spheroid behavior. Furthermore, we show that CAF-mediated collagen matrix remodeling and degradation precedes MCF7 spheroid disruption and is critical in promoting cancer cell spheroid expansion and cell dissemination. While broad-spectrum matrix metalloproteinase inhibition suppresses CAF-driven collagen degradation and MCF7 spheroid expansion, it does not prevent ECM remodeling, CAF migration, or single-cell dissemination of cancer cell spheroids. Furthermore, a complementary heterospheroid model reveals similar ECM remodeling and invasion dynamics despite altered cellular arrangement of cancer cells and CAFs. These findings enhance our understanding of the relationship between CAF activity and collagen matrix remodeling processes that promote cancer cell invasion, providing insights into the potential therapeutic benefits of targeting CAFs in breast cancer treatment.

## 1. Introduction

Breast cancer is one of the most commonly diagnosed cancers worldwide and comprises distinct molecular subtypes characterized by specific biomarkers and clinical features [1]. These subtypes arise from complex intra- and inter-cellular interactions driven by genetic mutations, epigenetic modifications, and dynamic crosstalk between cancer cells and their heterogeneous tumor microenvironment (TME) [2]. It has been established that Luminal A breast cancer – the most prevalent subtype - generally follows a more indolent course and has a favorable prognosis. However, a subset of Luminal A tumors associated with α-smooth muscle actin (α-SMA)-positive cancer-associated fibroblasts (CAFs) in the TME exhibits significantly aggressive behavior, leading to distant metastasis and relapse [3].

Within the TME, CAFs, along with other cellular components such as endothelial cells, immune cells, and the extracellular matrix (ECM), collectively constitute the tumor stroma. It has been shown that the stromal-to-tumor ratio serves as an independent prognostic indicator, with a higher stromal fraction correlating with increased tumor aggressiveness and poorer patient outcomes [4–6]. However, substantial variability across studies highlights the complexity of the stromal contributions to tumor progression [4–9]. These discrepancies likely stem from the dynamic and reciprocal interactions between cancer cells and their stromal counterparts, leading to extensive reorganization of cellular composition and ECM architecture during tumorigenesis [10,11]. Consequently, these processes create a heterogeneous TME characterized by spatially distinct features, underscoring the need for a more refined analytical framework to accurately assess the stromal contributions to disease progression. The inherent heterogeneity of the TME results in significant phenotypic diversity among cancer cells and CAFs, leading to unique functional signatures and cellular characteristics that vary across different regions of the TME [12].

In Luminal breast cancers, CAFs actively promote tumor progression by reprogramming cancer cells, which enhances therapeutic resistance, disrupts cancer cell metabolism, and increases cancer cell motility [13]. Direct interactions between CAFs and cancer cells further enhance tumor cell motility and metabolism, sustaining tumor progression [13,14]. CAFs also exert considerable regulatory effects on the collagen ECM through mechanisms that modulate collagen cross-linking, deposition, and degradation [15]. CAFs mediate collagen degradation through the secretion of matrix metalloproteinases (MMPs)[16]. In addition to this proteolytic activity, they also contribute to ECM remodeling by exerting contractile forces that realign collagen fibers [17–20]. This biomechanical remodeling leads to the aggregation of thin collagen fibrils into thicker, aligned bundles - referred to as “collagen highways” - which serve to facilitate and guide cancer cell migration and invasion[21].

As the TME undergoes temporal evolution, its ECM is gradually remodeled into a tumor-facilitating matrix due to the biophysical and biological activities of CAFs and cancer cells. These cells can modulate the biophysical and biochemical properties of the collagen ECM, leading to increased matrix stiffness, which is a notable characteristic of breast malignancy [22,23]. *In-vitro* models support these findings, showing that aligned collagen fibers increase matrix stiffness and improve tumor cell migration efficiency by promoting directional persistence [24]. These structural and mechanical changes to the ECM create a microenvironment that facilitates tumor progression and enhances therapeutic resistance [25]. Yet, the role of CAFs in indolent breast cancers, characterized by non-migratory cancer cells, and the mechanisms that may precipitate invasion and eventual metastasis in these settings remain unresolved. Specifically, the effects of CAF-induced matrix remodeling, along with its reciprocal influence on CAF–cancer cell interactions in indolent breast cancer, are not well defined. Recent advances in breast cancer research have emphasized that local ECM architecture at the TME boundary, along with the spatial organization of cancer cells and CAFs, significantly affects patient outcomes [26–30]. To address this, we studied the behaviour of luminal breast cancer spheroids in collagen matrix and ECM remodeling in the presence of breast cancer CAF spheroids. Our findings enhance the understanding of the relationship between CAF activity and collagen matrix remodeling processes that promote cancer cell invasion, providing insights into the potential therapeutic benefits of targeting CAFs in breast cancer treatment.

## 2. Materials & Methods

### 2.1. Cell culture and reagents

The human Luminal A epithelial breast cancer cell line MCF7 was purchased from ATCC. The human telomerase reverse transcriptase (hTERT)-immortalized breast 19TT (CAFs) cells have been previously described [31]. All cells were cultured in Dulbecco’s Modified Eagle Medium (DMEM), High Glucose (Sigma) containing 4.5 g L−1 glucose, l-glutamine but without sodium pyruvate and supplemented with 10% fetal bovine serum (FBS, Sigma) and 1% antibiotic–antimycotic solution (Gibco). Cells were frequently tested for the absence of mycoplasma and checked for authenticity by short tandem repeater (STR) profiling. For MMP inhibition experiments, spheroids were treated with Batimastat (BB94, 30 µM; Selleckchem) by incorporating the drug into the culture medium, following the manufacturer’s instructions. Vehicle controls were included accordingly.

### 2.2. Lentiviral Transduction

Constitutive expression of fluorescent markers was accomplished via lentiviral transduction. MCF7 cells were labeled with plv-mCherry, and 19TT cells were labeled with pLenti CMV H2B green fluorescent protein (GFP) Hygro (656-4). pLenti CMV GFP Hygro (656-4) was a gift from Eric Campeau & Paul Kaufman (Addgene plasmid #17446; http://n2t.net/addgene:17446; RRID:Addgene_17446). Lentiviruses were produced by co-transfecting cDNA expression plasmids with helper plasmids pCMV-VSVG, pMDLg-RRE (gag/pol), and pRSV-REV into HEK293T cells using polyethyleneimine (PEI). Cell supernatants were harvested 48 hours after transfection and stored at −80°C. Cells were labeled by infecting for 24 hours with respective lentiviral supernatants diluted 1:1 with cell culture medium and 5ng/ml of polybrene (Sigma-Aldrich). 48 hours after infection, 19TT cells were placed under Hygromycin-B (Thermo Fisher Scientific) selection, and MCF7 cells were placed under Neomycin (A1720, Sigma-Aldrich) selection. The fluorescently labeled cells were cultured with 100 µg/ml of respective antibiotics to obtain a stable fluorescent cell lines and maintain selection pressure.

### 2.3. Spheroid seeding and characterization

MCF7 and 19TT-CAF spheroids were seeded in Corning Elplasia 96-well Round Bottom Ultra-Low Attachment (ULA) Microcavity Microplates (Corning, 4442). Briefly, cells were trypsinized (Trypsin-EDTA, GibcoTM, 25200056) and resuspended to a concentration of 1×10^6^ cells/ml. 40 × 10^3^ (500 cells per microwell, 79 microwells) are seeded to produce 79 spheroids. Spheroids were ready after 48 hours of culture. MCF7 spheroids that had a size of 175 ± 25µm in diameter and 19TT-CAF spheroids that had a size of 125 ± 25µm in diameter were used for invasion studies. MCF7 spheroids were cultured in DMEM, High Glucose (Sigma) containing 4.5 g L−1 glucose and l-glutamine but without sodium pyruvate and supplemented with 10% fetal bovine serum (FBS, Sigma) and 1% antibiotic–antimycotic solution (Gibco). For 19TT-CAF spheroids, a high-viscosity culture medium was made using methylcellulose as described previously [32]. Briefly, a 20% methylcellulose solution was made by sterilizing 6g of methylcellulose in an autoclave. This was subsequently mixed with 250 ml of DMEM High Glucose. Once methylcellulose was dissolved, an additional 250ml of DMEM High Glucose and 55ml FBS was added, and the solution was mixed overnight at 4°C. The following morning, the solution was centrifuged at 5000g for 2h at 4°C, and the supernatant was stored at 4°C until use.

Heterogenous spheroids were generated using the hanging drop method. Briefly, cells were trypsinized (Trypsin-EDTA, GibcoTM, 25200056) and resuspended to a concentration of 1 × 10^3^ cells/ml. Cell lines were mixed and diluted to a final concentration of 1000 cells / 30µL, consisting of 500 cells of each cell line. As described above, a high-viscosity culture medium, made using 20% methylcellulose (M0512, Sigma), was used to culture cells. 30µL droplets containing 1000 cells were cultured 48 hours before the fully formed spheroids were harvested for experiments.

### 2.4. Preparation of collagen hydrogels

Collagen type-1 hydrogels were prepared on ice to prevent polymerization. Briefly, collagen (Corning™, 354249) was mixed with 10x Reconstitution Buffer (200 mM HEPES + 262 mM sodium bicarbonate) and 10x DMEM (Gibco™, 12100061) in an 8:1:1 ratio, adjusted to pH 7.4 using 0.5M NaOH, and diluted with sterile MiliQ water to the required concentration (3 mg/ml) [33]. The hydrogel was incubated at 4°C for 20 minutes to promote fiber formation, followed by 37°C for 40 minutes.

### 2.5. Spheroid Invasion Experiments

A thin 100µL collagen bed (3 mg/ml) was created and allowed to gel for the invasion experiment (as mentioned in section 2.4). This was followed by harvesting 19TT-CAF and MCF7 spheroids, mixing them with freshly prepared collagen, embedding them on top of the collagen bed, and incubating them as stated above. Culture media was added after incubation, and the device was set up for time-lapse imaging. Initial imaging for the first experiment was conducted 1 hour and 10 minutes following spheroid seeding (or immediately post collagen gelation). This time point was designated as *T*_0_ for all subsequent analyses in this study. The culture medium was refreshed every 48 hours for experiments running until 96 hours. Analysis was limited to spheroids (heterogenous or dual-spheroid systems) that were more than 400 µm apart from other spheroids at the beginning of the experiment. This ensured that no additional interactions would occur within the region of interest (ROI) for the experiment duration.

### 2.6. Time-lapse Confocal and Reflection Imaging

Confocal fluorescence and reflection microscopy were performed using the Zeiss LSM-980 integrated with a live cell incubator. Temperature was maintained at 37°C and CO_2_ at 5% during microscopy measurements. Images were collected at 2-hour intervals and analyzed using ImageJ2 (v2.14, National Institute of Health, USA) and custom made MATLAB scripts. For image acquisition, EC Plan-Neofluar 10x/0.3NA M27 (Zeiss) air objective was combined with 13mW 488 nm and 543 nm lasers for fluorescence imaging. During acquisition, the digital gain was set to 1, digital offset 0, laser power to 0.2%, and master gain at 650V. Appropriate band-pass filters were used to prevent spectral overlap between GFP and mCherry channels. A 488 nm laser with a detection range of 476 nm to 502 nm was used for reflection microscopy. The master gain was set to 600V, and 0.2% laser power was used during reflection imaging. The reflection images in the figures were collected with equivalent gain settings and presented with equivalent digital contrast, brightness, and levels. All images are acquired at an optical zoom of 1.0, with a field of view of 848.53μm x 848.53μm with 2048×2048 pixels. Z-stacks were acquired at 20μm step size for a total range of 400μm.

### 2.7. Image Processing

CZI files were extracted and saved as ‘<u>.tiff</u>’ files after image acquisition using custom ImageJ scripts. Before analysis, these files were processed, during which mean noise was subtracted from images, and fluorescent channels (GFP and mCherry) were checked for spectral overlap.

#### 2.7.1. Circularity and Change in Area of MCF7 Spheroids

MCF7 spheroid images were extracted and saved as image sequences using custom ImageJ scripts. Maximum Z-projections were created to find the maximum diameter of the MCF7 spheroids. These maximum projections were then utilized with custom MATLAB scripts to binarize and threshold images to identify MCF7 spheroid edges. In-built *regionprops* function was subsequently used to measure the change in area and circularity of spheroids. To calculate the shortest distance between MCF7 and 19TT CAF spheroids, image sequences in the Z-plane at *T*_!_ were saved for each spheroid. Weighted centroids in three-dimensional space were computed based on the maximum spheroid area at each Z-plane. The 3D centroid-to-centroid distance was then determined. Using a nearest-neighbor algorithm, the coordinates of the intersection between the centroid-connecting line and the spheroid boundary were identified. These boundary coordinates for both spheroids were then used to calculate the shortest inter-spheroid distance.

#### 2.7.2. Migration Analysis

CAF spheroid migration was calculated by measuring the maximum cross-sectional spheroid area at *T*_0_ (*A*_0_) and during its culture within the collagen matrix (*A*_*t*_). The invasive index was then defined as δ*A* = (*A*_*t*_/*A*_0_) − 1 to quantify the level of invasion for different spheroids.

#### 2.7.3. Collagen Fiber Orientation, Void Fraction and Brightness Characterization

Collagen fiber images obtained via reflection imaging were analyzed using ImageJ and MATLAB. Collagen planes corresponding to the central spheroid plane were selected in ImageJ for further analysis. Spheroid area and position are segmented from the image using Gaussian filtering (sigma = 2 pixels), morphological operations and Otsu’s thresholding [34]. OrientationJ’s vector field analysis is used to determine collagen fibers’ local orientation and isotropy with respect to the absolute x-axis [35]. The size of the gaussian kernel was set to 10 µm [35,36]. As shown in Supporting Fig. S1, the resulting orientation mask was analyzed in conjunction with intensity thresholded collagen masks using custom MATLAB scripts to identify, transform, and subsequently calculate the collagen fiber alignment relative to the centroids of the corresponding spheroids, providing a detailed spatial understanding of collagen architecture about spheroid positioning. To quantify collagen degradation, we extended on our MATLAB script and used the intensity thresholded collagen masks to identify void spaces between collagen fibers based on relative pixel intensity values and calculate the void fraction (*V*_*f*_) of the collagen plane. These *V*_*f*_ values were then normalized to the *V*_*f*_ value at *T*_0_. To quantify the change in collagen fiber brightness (pixel intensity values), we summed up all pixel values of the collagen fibers at each time point (*I*_*P*_) and normalized them by *T*_0_.

### 2.8. Tissue Sectioning, Imaging and Analysis

#### 2.8.1. Tissue Processing, Histochemical Staining, and Digitization of Human Breast Cancer

Formalin-fixed, paraffin-embedded tumor tissues were sectioned into four µm slides and processed sequentially, including deparaffinization with xylene, rehydration through an ethanol gradient, histological staining, air drying at room temperature, and mounting with a xylene-based medium (Permount; Thermo Fisher Scientific Inc., Waltham, Massachusetts, USA). For this study, a tissue block containing the most invasive tumor region, used for Tumor (T)-status assessment, was selected to prepare serial sections. These sections were stained with hematoxylin and eosin (H&E) and Picrosirius red (PSR). PSR staining involved treatment with 0.1% Direct Red 80 dye in saturated picric acid (Sigma-Aldrich, Merck, Darmstadt, Germany) for 60 minutes, followed by rinsing in two changes of 0.5% glacial acetic acid (Avantor, Inc., Radnor Township, Pennsylvania, USA). Stained slides were digitized in brightfield mode using a Pannoramic 250 slide scanner (3DHISTECH, Budapest, Hungary) at 200× magnification (0.39 µm per pixel) and saved as proprietary .mrxs files.

#### 2.8.2. Fluorescent Whole-Slide Imaging of Picrosirius Red-Stained Tissue Sections

Picrosirius red-stained tissue sections were digitized in fluorescent mode using a Pannoramic 250 slide scanner (3DHISTECH, Budapest, Hungary) at 200× magnification (0.39 µm per pixel) and saved as proprietary .mrxs files. Tissue autofluorescence was captured using a FITC filter cube (excitation: 460–488 nm, emission: 502–547 nm) with an exposure time of 800 m seconds. The fluorescence emission of the fibrous ECM stained by Sirius red was recorded using a TRITC filter cube (excitation: 532–554 nm, emission: 573–613 nm) with an exposure time of 44 ms.

#### 2.8.3. Immunohistochemistry

For multiplex immunofluorescence, tissue sections were deparaffinized, rehydrated, and subjected to heat-induced epitope retrieval using citrate buffer (pH 6.0). To minimize non-specific binding, sections were incubated with 5% goat serum for 10 minutes at room temperature. Primary antibodies were applied overnight, including anti-αSMA (clone 1A4, DAKO, M0851, 1:300 dilution) and pan-cytokeratin (AE1/AE3, DAKO, M3515, 1:300 dilution). The next day, sections were incubated with Alexa Fluor-conjugated secondary antibodies: 647 goat anti-mouse IgG2a (Life Technologies, A21241, 1:200 dilution) and 546 goat anti-mouse IgG1 (Invitrogen, A21123, 1:200 dilution). All antibody incubations were performed for 30 minutes at room temperature in the dark. Finally, nuclei were counterstained with Hoechst 34580 (Sigma-Aldrich, 63493, 1:2000 dilution) for 5 minutes.

### 2.9. Statistical Analysis

All graphs shown in this work were produced using GraphPad Prism. Statistical analyses were run within the software, and the two groups’ significance was measured using unpaired t-tests or ANOVA based on experiment variable dependencies. Statistical significance between more than two groups was assessed using ANOVA tests. Probability values less than 0.05 were deemed significant (p < 0.05 *, p < 0.01 **, p < 0.001 ***).

## 3. Results

### 3.1. Cancer cell and CAF distribution, and collagen morphology observed in luminal breast tumor sections

To explore the heterogeneity of the TME and the organization of the ECM in indolent breast cancers, we analyzed tissue sections from Luminal A breast cancer samples. Our focus was on the collagen architecture and cellular organization at distinct tumor boundaries (Fig. 1 A-C). The analysis of multiple tumor boundaries across four samples revealed significant variability in the collagen fiber organization in the TME ECM. Some tumor boundaries displayed a thick collagen layer that separated the TME from adjacent healthy tissues, while others were characterized by infiltrative collagen fiber projections extending into the surrounding tissue. Based on these structural differences, we classified tumor boundaries as either non-infiltrative or infiltrative tumor boundaries (Fig. 1B, C). Among the four tumor samples analyzed, two (Tumor ID: CAF01, CAF02) exhibited both infiltrative and non-infiltrative tumor boundaries, while the remaining two (CAF03, CAF06) displayed only one type. Tumor section CAF03 displayed exclusively non-infiltrative margins, with most CAFs located distinctly away from tumor cell clusters. In contrast, CAF06 exhibited infiltrative boundaries, with CAFs interspersed among tumor cells. The tumor shown in Fig. 1 (Tumor ID: CAF01) featured both infiltrative and non-infiltrative fronts, while the remaining sections are presented in Fig. S2. Infiltrative tumor fronts were marked by highly organized collagen fibers running perpendicularly to the tumor boundary and forming protrusions that extended into adjacent mammary fat tissue (Fig 1A, blue arrows). In contrast, non-infiltrative fronts exhibited less organized collagen running parallel to the tumor boundary, thereby creating a distinct and continuous boundary between the tumor and surrounding tissues (Fig 1A, red arrows). Immunohistochemical staining for pan-Cytokeratin (pCK), an epithelial cell marker used to identify cancer cells within the tumor boundary, and α-SMA, a marker for activated fibroblasts used for CAFs, revealed observable differences in cellular composition between regions. At these infiltrative fronts, cancer cells were organized into small clusters interspersed with CAFs, as reflected by the proximity of pCK and α-SMA signals. Additionally, infiltrative tumor boundaries demonstrated a pronounced localization of α-SMA-positive cells at the tumor–tissue interface. In contrast, non-infiltrative boundaries exhibited a more distal distribution of α-SMA expression relative to the tumor margin. These findings highlight the spatial heterogeneity of the stroma and its critical role in remodeling the ECM at infiltrative tumor margins. They also reveal the complex interplay among cancer cells, CAFs, and collagen organization in modulating breast cancer invasiveness.

**Figure 1.**
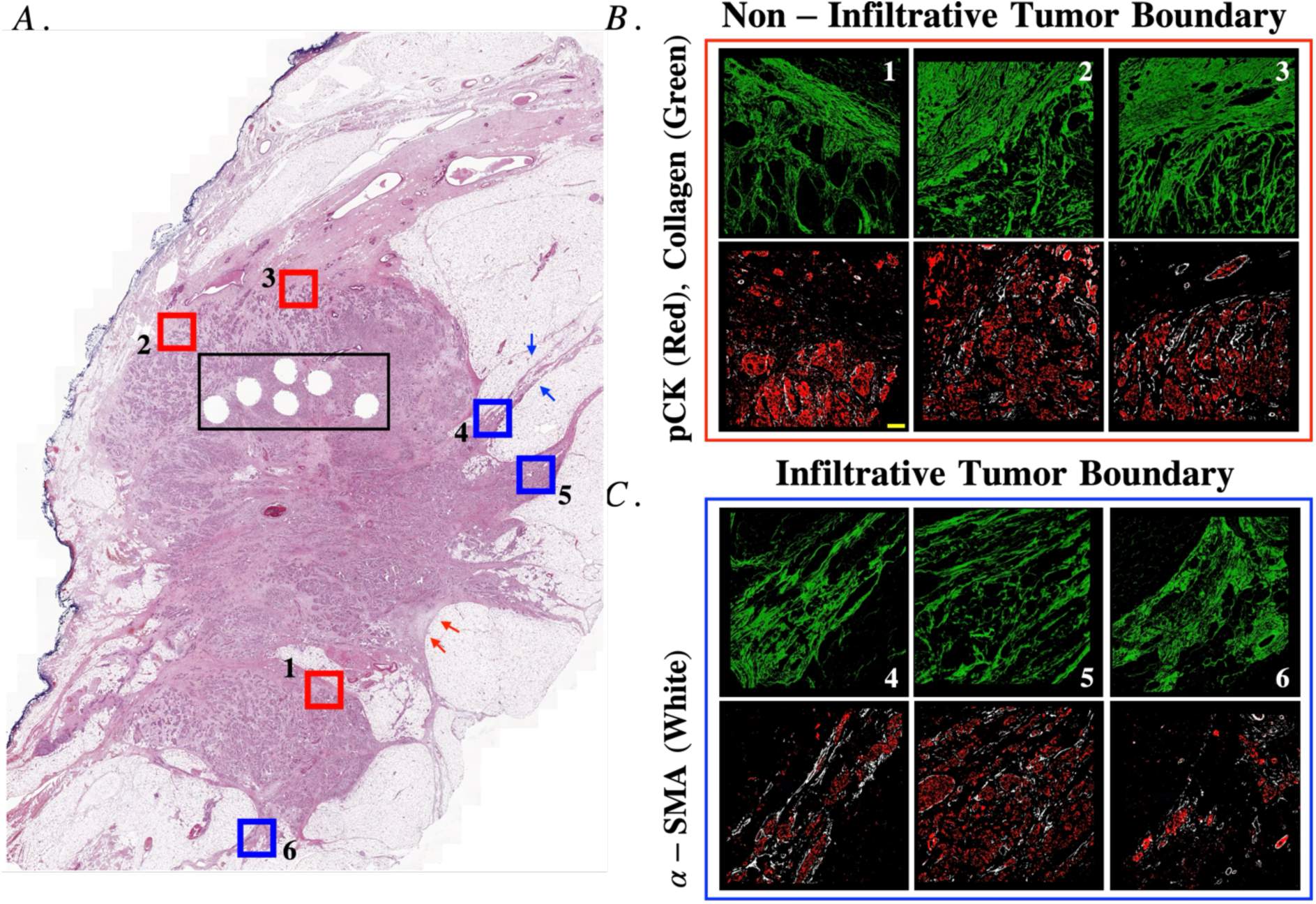
Infiltrative and Non-Infiltrative TME boundaries can be identified based on cellular organization and collagen architecture. A. H&E Stained tissued section of Luminal A tumor. Holes visible at the centre of the tumor (black box) are biopsy sections. ROIs boxed in blue and red are indicative of infiltrative and non-infiltrative tumor boundaries respectively. Blue arrows indicate infiltrative tumor boundary with collagen fingers protruding into the breast tissue. Red arrows indicate non-infiltrative tumor boundary creating a distinct boundary between the tumor core and breast tissue. B. Fluorescent scans of Non-Infiltrative tumor boundaries. On top (red) is the collagen organization at the tumor boundary and below is the scanned sequential tissues slides indicating a-SMA (CAF, white) and pCK (tumor cells, red) organization at the tumor boundary. Scale Bar = 100 µm. C. Fluorescent scans of Invasive tumor boundaries. On top (red) is the collagen organization at the tumor boundary and below is the scanned sequential tissues slides indicating ⍺-SMA (CAF, white) and pCK (tumor cells, red) organization at the tumor boundary.

Despite these qualitative insights, several fundamental questions remain. A key challenge is understanding how spatio-temporal modifications of the collagen ECM induced by CAFs affect the structural integrity and dissemination of cancer cells. This evaluation is complex and cannot be directly assessed from tissue samples alone. Addressing this challenge requires differentiating between the CAF-induced mechanical pulling of ECM and the enzymatic degradation that occurs during collagen remodeling to clarify their distinct impacts on CAF and cancer cell behavior. This underscores the need for a dynamic *in-vitro* assay capable of capturing real-time interactions between CAFs, the ECM, and cancer cells. Furthermore, the role of CAF spatial distribution relative to tumor spheroids in determining the pattern and extent of ECM remodeling, as well as cancer cell dissemination, remains unclear. To unravel these biophysical mechanisms underlying cancer invasion, it is required to establish an *in-vitro* cancer model that allows for time-resolved analysis of CAF-mediated ECM remodeling and tumor cell dynamics.

### 3.2. CAF mediates ECM Remodeling and the subsequent MCF7 Spheroid Expansion and Dissemination

To investigate the impact of distinct cellular organization at infiltrative and non-infiltrative tumor fronts on intercellular interactions and ECM remodeling, we co-embedded 19TT-CAF and MCF7 cancer cell spheroids within a 3D collagen matrix (3 mg/ml). This assay configuration enabled us to analyze key features observed in tissue sections, such as the interspersed cellular architecture, orientation of collagen fibers, and loose tumor clusters that could interact with 19TT-CAF cells, while allowing for kinetic visualization of ECM remodeling (See Fig 2A, B, D). In this co-culture system, 19TT-CAF spheroids maintained direct access to the collagen matrix and were able to engage with MCF7 tumor spheroids through short-distance migration. Fig. 2(B) and movies 1, 2 and 3 illustrate the spatial organization between 19TT-CAF and MCF7 spheroids compared to a single MCF7 spheroid embedded in the collagen matrix. Single MCF7 spheroids displayed gradual nearly uniform expansion over the entire 48-hour period. However, when 19TT-CAF spheroids were seeded less than 200µm apart from MCF7 spheroids, we observed non-uniform expansion and eventual collapse of MCF7 spheroids. In several samples, we also observed single-cell dissemination from MCF7 spheroids (Fig 1D, MCF7, 48 hours, *d* < *D_c_*). Conversely, when the inter-spheroid distances (*d*) were greater than 200µm, MCF7 spheroids exhibited significantly less, yet non-uniform expansion after 48 hours. We quantified the effect of 19TT-CAF and MCF7 inter-spheroid distance on MCF7 spheroid expansion using the maximum diameter of MCF7 spheroids (*D_c_* = 200µ*m*) as a reference and comparing spheroid expansion when 19TT-CAF spheroids were seeded at distances *d* < *D_c_* and *d* > *D_c_*. To estimate the overall expansion of MCF7 spheroids, we calculated the projected spheroid area at *T*_0_ and after 48 hours. The expansion of the spheroid relative to its initial area was then calculated as *dA* = (*A*_48_ – *A*_0_)/*A*_0_. As shown in Fig 2(C), for MCF7 spheroids embedded alone in the collagen matrix, we observed a change in the spheroid area of 1.1 after 48 hours. For 19TT-CAF spheroids seeded at a distance *d* < *D_c_*, the change in MCF7 spheroid area was 4.1 compared to 1.6 when 19TT-CAF spheroids separated by distances *d* > *D_c_*.

**Figure 2.**
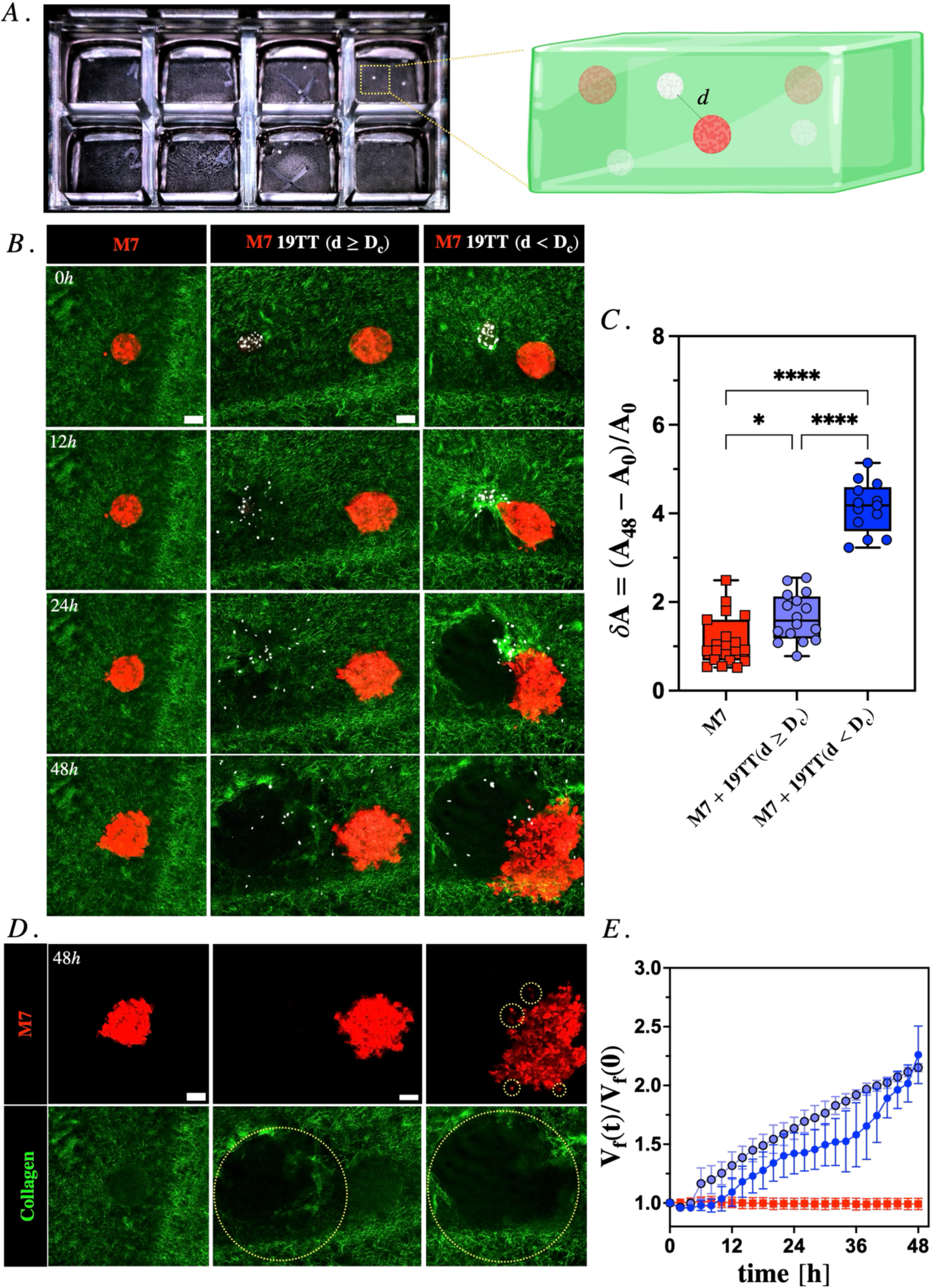
19TT CAF - Mediated Disruption of MCF7 (M7) Spheroids is Dependent on Inter-Spheroid Distance. (A) Ibidi 8-well glass-bottom slide with a 3D collagen matrix containing multicellular tumor spheroids. A zoomed-in schematic represents multicellular luminal M7 breast cancer and CAF spheroids embedded in a 3D collagen matrix, separated by a distance “d.“ (B) Representative fluorescence images of M7 mCherry-labeled cancer cell spheroids (red) and 19tt H2B GFP-labeled CAF spheroids (white) taken at 0 h, 24 h, and 48 h post-embedding in a 3 mg/ml rat-tail collagen matrix (green). Spheroids were separated by distances of (ii) 450 µm and (iii) 150 µm. (C) Quantification of the M7 (MCF7) spheroid area was performed under three conditions: when embedded alone (n = 9), with 19TT spheroids within 200 µm (n = 9), and with 19TT spheroids at distances greater than 200 µm (n = 9) after 48 h. M7 spheroids seeded alone only doubled in size. Tumor spheroids seeded more than 200 µm away exhibited a 2.5-fold increase. In comparison, tumor spheroids seeded less than 200 µm from 19TT CAF spheroids demonstrated a nearly four-fold increase in area over 48 h (*p < 0.05, ****p < 0.0001). (D) Overview of (1) single-cell dissemination (yellow circles) observed from M7 spheroids and (2) collagen matrix integrity (degraded collagen area in yellow circles) under three conditions: (i) M7 alone, (ii) M7 + CAF, *d* ≥ *D_C_*, and (iii) M7 + CAF, *d* < *D_C_*, after 48 hours. (E) Void Fraction (*V_f(t)_*/*V_f_*_(0)_) Quantification of the collagen hydrogel measured from 0 to 48 h and normalzied to its initial value *V_f_*_(0)_ for M7 (red) (n = 9), M7 + CAF, *d* ≥ *D_C_* (n = 9), (dark blue) and (iii) M7 + CAF, *d* < *D_C_* (light blue), after 48 h (n = 9).

Based on the dynamics captured in Figure 2B and movies 1,2 and 3, we identified key time dependent interaction patterns involving 19TT-CAFs with both the ECM and MCF7 spheroids. Initially, we observed localized remodeling of collagen fibers in the vicinity of 19TT-CAF spheroids, indicating early mechanical engagement with the surrounding ECM. During the initial 12 hours, 19TT-CAF spheroids remodeled thin collagen fibres into thicker, radially aligned fibers. Image analysis across multiple Z-planes revealed that 19TT-CAFs drew collagen fibres from various focal planes, bundling them into radially aligned fibers within the central plane of the spheroids (Fig. S3). This process likely contributed to increased collagen fiber pixel intensities and deformation of the MCF7 spheroids, as shown in Fig. 2B (*d* < *D_c_*) [36,37]. Following initial collagen matrix remodeling, 19TT-CAF cells initiated migration into the surrounding remodeled collagen matrix. This invasion by 19TT-CAF cells was characterized by progressive matrix degradation, void formation and eventual dissemination of the MCF7 spheroids. To quantify matrix degradation, we calculated the void fraction (*V_f_*) of the collagen matrix through image analysis, normalizing it to the initial time point. The time evolution graphs in Fig. 2E indicated no collagen degradation for single MCF7 spheroids. In contrast, matrix degradation began to increase beyond t = 6 hours, demonstrating significant collagen degradation for both MCF7 + 19TT-CAF co-culture spheroid cases, which doubled its initial value by the end of 48 hours. 19TT-CAF spheroids at *d* < *D_c_* showed greater variability in collagen degradation rates as seen from the *V*_*f*_graph in Fig. 1E compared to 19TT-CAF spheroids at *d* > *D_c_*. The observed variability in collagen degradation rates could be explained by the spatial proximity between 19TT-CAF and MCF7 spheroids. At shorter distances, early interactions between the two cell types could have attenuated 19TT-CAF-mediated collagen remodeling. In contrast, when seeded at greater distances, 19TT-CAF spheroids would have been able to remodel the collagen matrix without such interference prior to engaging with MCF7 spheroids.

Interactions between migrating 19TT-CAF cells and MCF7 spheroids were characterized by the expansion of the MCF7 spheroid, followed by its eventual collapse into the void created within the collagen matrix. As shown in Fig 2(D), several disseminated MCF7 cells were observed for cases where *d* < *D_c_*. This observation could not be extended to MCF7 spheroids or MCF7 separated by *d* > *D_c_* from 19TT-CAF. These results confirm that inter-spheroid distance is a critical factor influencing MCF7 spheroid expansion, single-cell dissemination, and the rate of collagen matrix degradation. In the following sections, we will focus our analyses on the 19TT-CAF and MCF7 spheroid system where *d* < *D_c_*.

### 3.3. 19TT-CAF spheroid dissemination mediates collagen matrix remodeling and degradation

During 19TT CAF-mediated collagen remodeling, collagen fibers reoriented locally to align radially toward the center of the 19TT-CAF spheroid (Fig. 3A, Movie 4). To quantify this alignment, we identified thick collagen fibers and measured their angles relative to the spheroid center, as illustrated in Fig. S1. An angle approaching 0° referred as a higher degree of fiber alignment to the center of spheroids. Over the first 10 hours, collagen fiber alignment gradually intensified (Fig. 3A, 3B), with a pronounced peak observed at 0° by the end of the 10^th^ hour. Additionally, we noted an increase in collagen fiber brightness under consistent imaging conditions while the 19TT-CAF spheroids remodeled the collagen matrix. To quantify this remodeling, we measured collagen fiber pixel intensity through image analysis and normalized it to the pixel intensity observed at initial time. The analysis (Fig. 3C) revealed a consistent increase in pixel intensity from hours ∼ 0 to 12, indicating that collagen fiber reorientation and bundling were associated with this increase in brightness. After the 12-hour mark, we observed a slight increase in the variability of collagen pixel intensities. The dissemination of 19TT-CAF single cells occurred after the collagen fiber remodeling, suggesting that the restructured collagen matrix facilitated 19TT-CAF migration and invasion. During initial 19TT-CAF mediated collagen fiber reorientation, minimal matrix degradation was observed. This was further supported by the stable values of *V*_*f*_ during the initial 12 hours, indicating a consistent phase of matrix remodeling without significant degradation. In contrast, at later time points, the rate of collagen degradation increased, as evidenced by a steeper slope in the degradation curve (Fig. 3D). Collagen fiber orientation in the ECM surrounding MCF7 spheroids remained unchanged over 24 hours, indicating the absence of ECM remodeling (Movie 5, Fig 3D).

**Figure 3.**
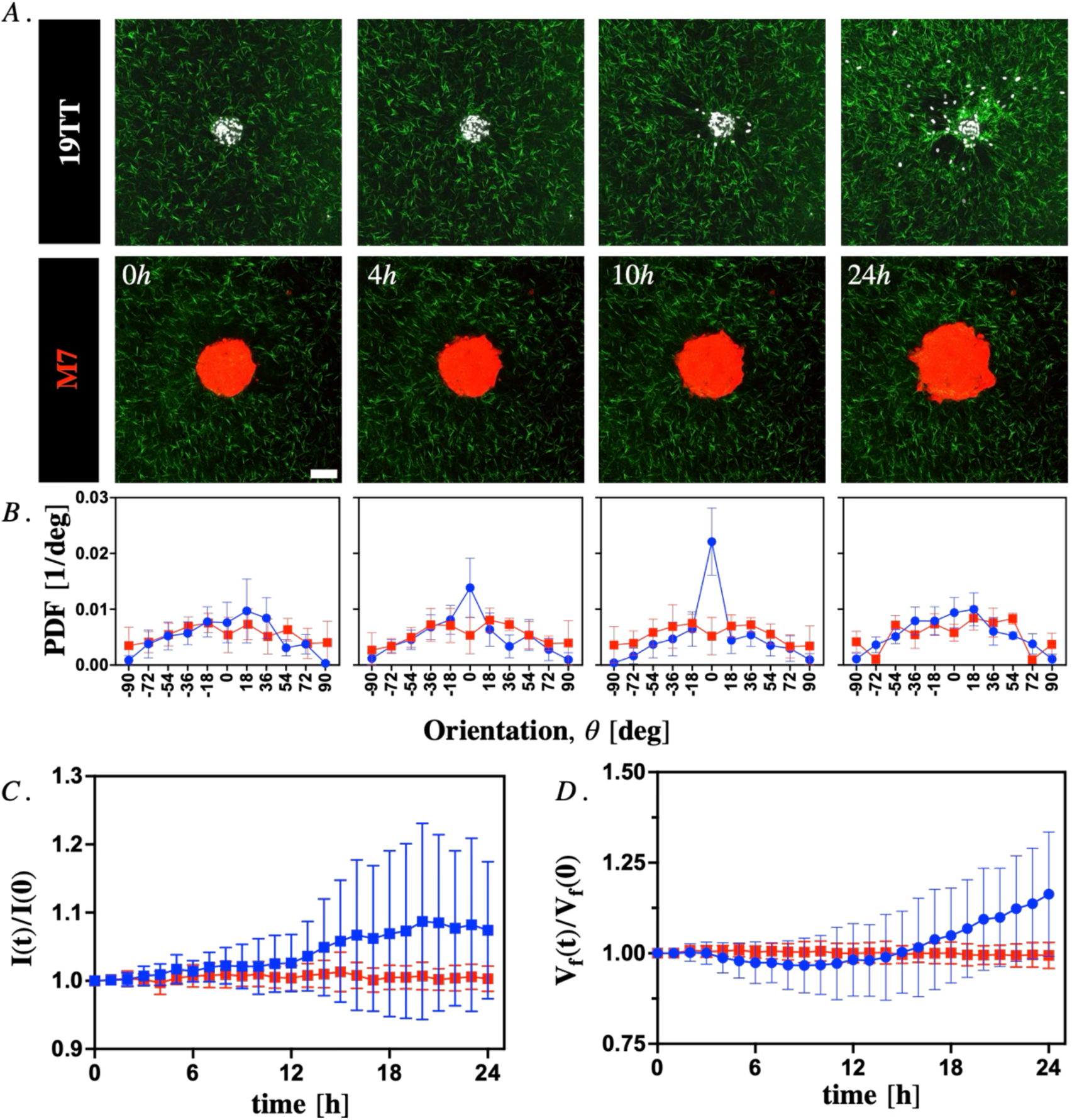
CAF spheroid-mediated matrix degradation is preceded by matrix remodeling and fiber reorientation. A. Representative time-lapse images of 19TT CAF (white) and M7 tumor spheroid (red) in a collagen matrix (green). B. Relative collagen fiber orientation concerning the CAF spheroid center in blue (n=11) and M7 spheroid center in red (n=10). Distribution of orientations from −90° to +90° is plotted on the x-axis and the probability density function (PDF) of orientations on the y-axis. Collagen fiber orientation sees a gradual increase in orientation to peak at 0° at the 10 h time point before the CAF spheroid begins disseminating. For M7 spheroids, the collagen fiber orientation remains relatively unchanged over 24 h. Scale bar = 100 µm. C. Quantification of collagen &_*t*_during the first 24 h of the experiment. M7 &_*t*_(red) remains relatively unchanged, but CAF &_*t*_ (blue) increases owing to collagen degradation. D. Quantified collagen fiber pixel intensities during the first 24 h of the experiment. Pixel intensities of collagen fibers surrounding CAF and M7 tumor spheroid-embedded collagen matrices are compared.

### 3.4. MMP Inhibition inhibits CAF-mediated matrix degradation but not CAF migration

Next, we investigated the influence of collagen matrix degradation on 19TT-CAF invasion and MCF7 spheroid expansion. To inhibit MMP-mediated matrix degradation we used the broad-spectrum MMP inhibitor Batimastat (BB-94) [38]. In comparison to the control condition, 19TT-CAF spheroids treated with BB-94 did not degrade the collagen matrix, resulting in no significant change in the void within the matrix over 24 hours (Fig. 4A, BB94 – Movie 6, Veh Control – Movie 7). Further quantification confirmed this observation: while *V_f_* increased in control conditions, it remained nearly unchanged for CAF spheroids treated with BB-94. However, BB-94 did not prevent collagen remodeling, as we observed thick collagen fibers at 10 hours in both the vehicle control and BB-94 conditions. Quantifying the orientation of collagen fibers showed no substantial differences between the vehicle control and BB-94 conditions (Fig. 4B). Additionally, lack of collagen matrix degradation resulted in increased pixel intensities over 24 hours (Fig. 4D) compared to control conditions, indicating that BB-94 did not impair the 19TT-CAFs’ ability physically remodel the collagen matrix. To assess whether collagen degradation influenced 19TT-CAF motility, we calculated the invasive index (*I_v_* = *A_t_*/*A*_0_ − 1), defined as the increase in spheroid cross-sectional area (*A_t_*), to quantify migration dynamics. For 19TT-CAF spheroids we found no significant difference in *I_v_* between control and BB-94-treated spheroids (Fig. 4D). These findings suggest that MMP inhibition had a minimal impact on 19TT-CAF invasion and single-cell dissemination. As shown in Fig. S4, MCF7 spheroids demonstrated no matrix remodeling activity under either control or MMP inhibition conditions. These results confirm that MMP activity is responsible for 19TT-CAF-induced matrix degradation but is not essential for 19TT-CAF-mediated matrix remodeling and subsequent 19TT-CAF collagen invasion.

**Figure 4.**
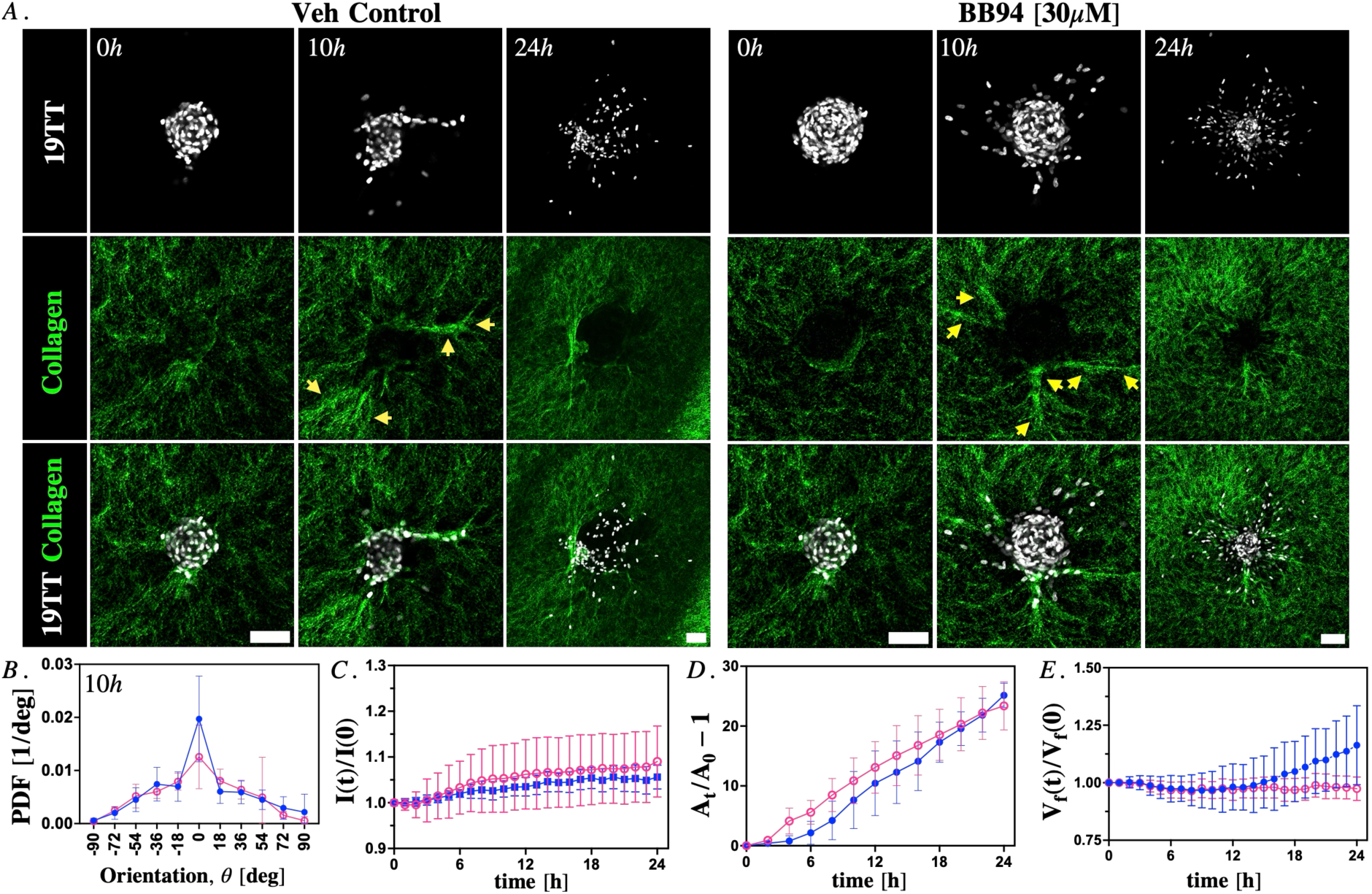
Broad-spectrum MMP with Batimastat (BB-94, 30µM) inhibition attenuates collagen matrix degradation but fails to suppress cancer-associated fibroblast (CAF) invasion or ECM remodeling. A. Representative images of CAF spheroids (white), the corresponding collagen matrix (green), and composite overlays for vehicle control and broad-spectrum MMP inhibition (Batimastat, 30 µM) conditions at 0 h and 10 h. A zoomed-out overview at 24 h highlights migrating CAF cells. Yellow arrows indicate reoriented collagen fibers in vehicle control and MMP inhibition conditions. B. Probability density function (PDF) of relative collagen fiber orientation concerning the CAF spheroid center for vehicle control (blue) and MMP inhibition (pink) conditions after 10h. C. Collagen fiber pixel intensities over 24 h. BB94 and vehicle control conditions exhibit a similar increase in collagen fiber pixel intensities over 24 h indicating broad-spectrum MMP inhibition does not impair CAF ability to remodel the matrix via application of contractile forces. D. The invasive index of CAF spheroids was observed 24 h post-embedding in the collagen matrix under vehicle control (n = 8) and MMP inhibition (n = 8) conditions. CAF migration dynamics remain similar across both conditions. E. Normalized collagen &_*t*_ comparing vehicle control and MMP inhibition conditions. Broad-spectrum MMP inhibition prevents collagen degradation, resulting in no significant change in collagen void fraction over 24 h.

### 3.5. MMP inhibition reduces MCF7 spheroid expansion but cannot prevent MCF7 cell dissemination

After evaluating MMP inhibition in 19TT-CAF spheroids, we expanded our analysis to the MCF7-19TT-CAF system. Time-lapse imaging showed that MMP inhibition significantly reduced the expansion of MCF7 spheroids and prevented their disruption at the 48-hour mark compared to the vehicle control condition (Fig. 5A, 5B, Movie <u>8</u>, <u>9</u>). However, MMP inhibition led to an observable increase in single-cell dissemination from MCF7 spheroids within the first 48 hours of the experiment (Fig. 5C). Despite the 19TT-CAFs maintaining their ability to remodel the collagen matrix while treated with BB-94, MCF7 spheroids did not display the deformations observed in the vehicle control conditions. As can be seen in Fig 5C, the patterns of MCF7 spheroid expansion varied between the two treatments: in the vehicle control condition, the bulk of the MCF7 spheroid was deformed by the 19TT-CAFs through matrix remoedeling and subsequent degradation before single-cell dissemination and cancer cell spheroid expansion occurred. In contrast, under MMP inhibition, MCF7 spheroids released single cells in all directions without prior physical disruption from the 19TT-CAF spheroids during the experimental timeframe (48 hours).

**Figure 5.**
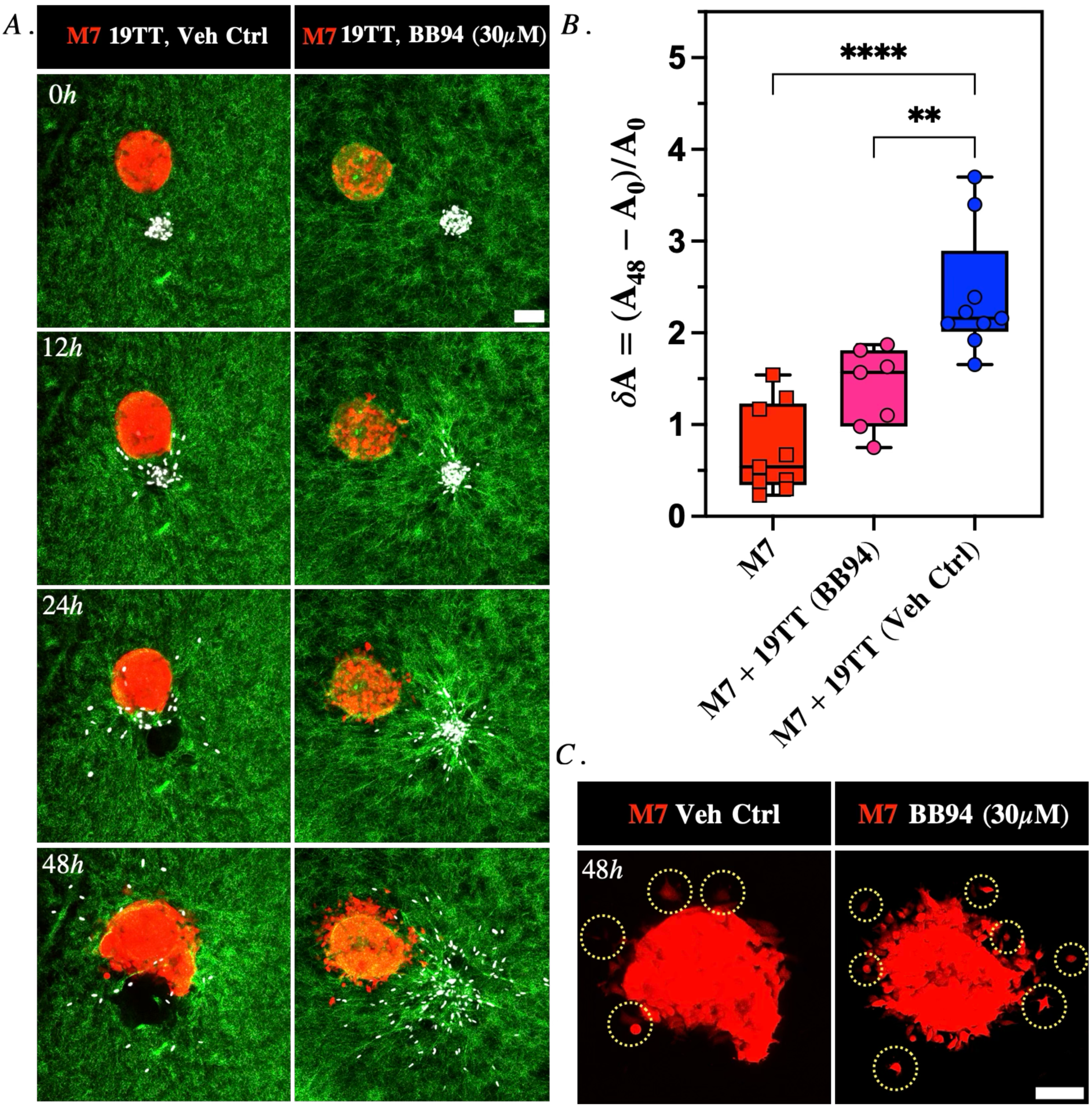
Broad spectrum MMP Inhibition with Batimastat (BB-94, 30µM) of the CAF-M7 spheroid system reduces M7 spheroid expansion but does not prevent single-cell dissemination. A. Representative time-lapse images of caf spheroids (white) and M7 tumor spheroids (red) embedded in a collagen matrix (green) in vehicle control and MMP inhibition (batimastat, 10µM) conditions. B. Quantification of M7 spheroid area changes when seeded alone (n=10), vehicle control (n=8) and MMP inhibition conditions after 48 h (n=6). C. Representative images of M7 spheroids in vehicle control and broad spectrum map inhibition conditions 48 h post embedding in a collagen matrix. Yellow arrows highlight single cells disseminating from the spheroids.

### 3.6. 19TT-CAF-MCF7 Heterogeneous spheroids display similar outcomes as dual-spheroid systems

To compare the results obtained from the 19TT-CAF-MCF7 dual-spheroid system with an alternative cellular organization, we cultured heterogeneous spheroids composed of approximately 500 MCF7 cells and 500 19TT-CAF cells each. The heterogeneous spheroids consistently exhibited a spatial architecture in which 19TT-CAF cells formed a central core surrounded by aggregating MCF7 cells (Fig. 6A). After embedding in collagen matrices, we conducted time-lapse imaging experiments using a protocol similar to that of the dual-spheroid system (Fig. 6A, Movie 10, <u>11</u>). Within the first 24 hours, 19TT-CAF cells migrated outward from the core of the hetero-spheroid into the surrounding collagen matrix in both control and BB94 conditions. Quantifying the relative orientation of collagen fibers in relation to the spheroid center (Fig. 6C) revealed comparable trends between the control and BB94 conditions. Initially, the orientation of collagen fibers was highly random; however, by the 24-hour mark, 19TT-CAFs within the hetero-spheroid had significantly reoriented the fibers, aligning them radially outward from the boundary of the spheroid. The expansion of the hetero-spheroid over the first 48 hours (Fig. 6D) mirrored the behavior observed in the dual-spheroid system, although we noted a greater number of single disseminating cells. Importantly, broad-spectrum MMP inhibition produced results consistent with those observed in the dual-spheroid system, significantly restricting spheroid expansion without affecting collagen fiber reorientation or the dissemination of MCF7 single cells.

**Figure 6.**
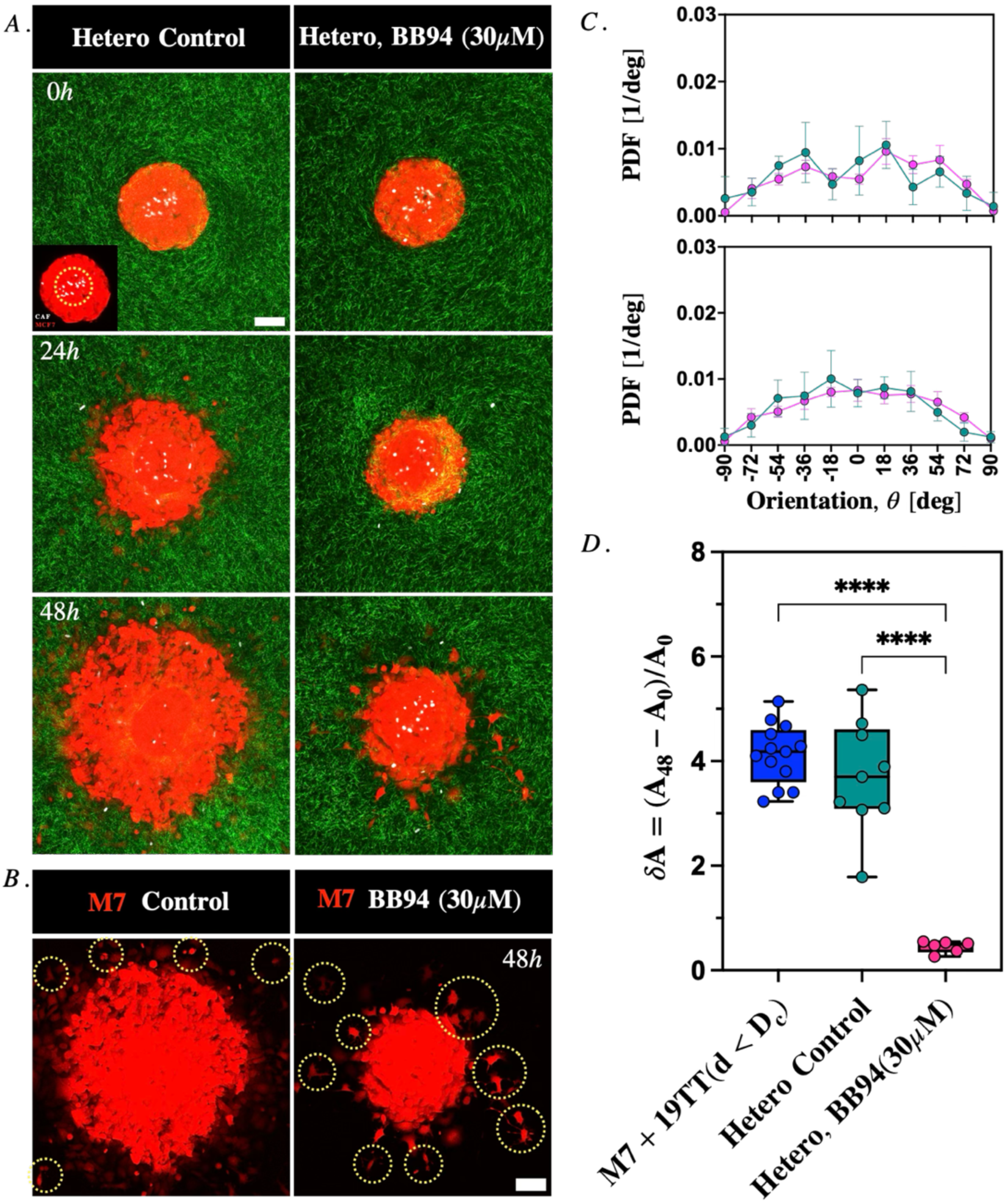
Heterospheroids (CAF – M7) show similar expansion rates and single cell dissemination as M7 spheroids when seeded close to CAF spheroids. A. Representative time-lapse images of caf and M7 tumor cells embedded as heterogeneous spheroids (red and white) in a collagen matrix (green) in vehicle control and broad spectrum map inhibition conditions (BB94, 30µM). Inset: Zoomed in Heterogenous Spheroid, yellow circle highights 19TT-CAF cells positioned at the centre of the spheroid. Scale bar = 100µm. B. Representative images of M7 tumor cells in vehicle control and broad spectrum map inhibition conditions 48 h post embedding in the collagen matrix. Yellow arrows highlight single cells disseminating from the spheroids. C. Collagen fiber orientation with respect to heterogenous spheroid center at 0 h and 24 h, respectively. Control in green (n=9) and MMP inhibition in pink (n=6) D. Quantifying heterogeneous spheroid area changes under vehicle control (n=9) and MMP inhibition (n=6) conditions after 48 h.

## 4. Discussion

The TME is a dynamic and heterogeneous setting that plays a critical role in tumor progression and invasion. Among its key components, CAFs significantly influence tumor development through various mechanisms, including ECM remodeling and cancer cell reprogramming. This study provides a systematic analysis of CAF-driven ECM remodeling and its impact on tumor spheroid expansion and single-cell dissemination. We began by characterizing the infiltrative and non-infiltrative tumor fronts in breast cancer tissue explants to better understand these interactions. Histological analysis of the infiltrative tumor fronts revealed a pronounced alignment of collagen fibers, elevated levels of α-SMA, and small clusters of cancer cells interspersed among CAF clusters. This indicates a highly interactive cancer cell-CAF landscape. In contrast, non-infiltrative tumor fronts exhibited larger, isolated tumor clusters with diminished collagen alignment and lower α-SMA expression, suggesting a less interactive and more segregated microenvironment.

Next, we aimed to capture the key features of the heterogeneous cellular architecture observed *in-vitro*, allowing for a systematic examination of interactions between CAFs and cancer cells, as well as between CAFs and collagen. The activity of 19TT-CAFs in the collagen matrix follows a dynamic sequence, beginning with structural reorganization of collagen fibers, followed by 19TT-CAF migration, and culminating in matrix degradation. Initial collagen matrix remodeling occurred concurrently with collagen polymerization, as evidenced by a transient increase in collagen fiber alignment at approximately 0° relative to the 19TT-CAF spheroid center, observable from the earliest imaging time points. This indicates that the collagen ECM was already undergoing remodeling by the time imaging began. During the first stage of matrix reorganization, 19TT-CAF spheroids bundled and aligned fibrillar collagen into thicker fibers, thereby preparing the matrix for subsequent invasion and degradation. The contractile forces exerted by 19TT-CAFs likely contributed to this effect by drawing collagen fibers into the imaging plane, enhancing their brightness, as demonstrated in Fig. S2. This brightness enhancement is the result of contractile forces applied by CAFs that lead to changing fiber orientation relative to imaging lasers, which is supported by reflection imaging studies showing that fiber alignment, in addition to fiber density, is one of the determinants of pixel intensity [36,37]. Following collagen reorganization, 19TT-CAF migration commenced and led to ECM degradation, characterized by a progressive increase in collagen over the subsequent 24 hours. However, the degradation rate plateaued between 24 and 36 hours, likely coinciding with 19TT-CAF-MCF7 interactions. After 36 hours, the degradation reaccelerated as 19TT-CAFs migrated deeper into the matrix, having already disrupted the MCF7 cancer cell spheroids. These observations highlight the spatiotemporal adaptability of 19TT-CAFs in ECM remodeling to facilitate tumor invasion. Proximity of 19TT-CAFs to MCF7 spheroids was a critical determinant of tumor expansion and dissemination, with spheroids located within one *D*_*C*_ of a 19TT-CAF spheroid experiencing significantly increased expansion and single-cell dissemination compared to those positioned further away.

To inhibit matrix degradation we employed broad-spectrum MMP inhibitor. MMP inhibition effectively prevented collagen degradation without impairing 19TT-CAF motility or ECM remodeling. Under these conditions, MCF7 cancer cell spheroid expansion was significantly reduced; however, single-MCF7 cell dissemination persisted and, in some cases, was enhanced compared to control conditions. This observation indicated that MCF7 spheroid expansion was dependent on the structural integrity of the collagen matrix, whereas the dissemination of individual cancer cells from the spheroid occurred independently of matrix integrity. Since MMP inhibition did not hinder CAF-driven matrix remodeling, we hypothesized that the contractile forces exerted by CAFs during collagen reorientation may create interstitial spaces between fibers, facilitating single-cell dissemination [36,39]. Analysis confirmed the absence of matrix degradation in MMP-inhibited samples. Evidence indicates that the architecture of the ECM plays a critical role in determining cell migration modes [40–42]. Denser matrices are associated with the dissemination and invasion of individual cells, whereas lower-density matrices tend to support and facilitate collective invasion [39,43,44]. These findings support the conclusion that ECM remodeling and degradation serve distinct functional roles: remodeling facilitates single-MCF7 cell dissemination, while degradation primarily supports bulk tumor expansion. Overall, these findings highlight the temporal sequence of events underlying CAF-mediated disruption of MCF7 spheroids. In a dual-spheroid system, CAF spheroids initiated matrix remodeling immediately upon embedding in the collagen hydrogel, as indicated by increased collagen fiber brightness and deformation of adjacent MCF7 spheroids. This process was followed by CAF invasion and subsequent collagen matrix degradation, as evidenced by the increase within the first 24 hours. The disruption of MCF7 spheroids occurred sequentially, leading to spheroid expansion and single-cell dissemination.

## 5. Conclusion

This study elucidates the spatiotemporal dynamics of CAF-driven ECM remodeling and its dual role in tumor invasion (focused on indolent breast cancer type), revealing valuable mechanistic insights into the interplay between stromal activity and tumor progression. Our findings demonstrate that CAFs orchestrate a sequential cascade of ECM reorganization, migration, and degradation, which collectively enable tumor spheroid disruption and dissemination. The initial phase of collagen fiber alignment and bundling by CAFs creates a permissive microenvironment for subsequent invasion, while MMP-mediated degradation facilitates bulk tumor expansion. Notably, MMP inhibition experiments revealed that ECM remodeling alone—via CAF contractility—suffices to generate interstitial spaces for single-cell dissemination, decoupling this process from matrix degradation. These results may also shed some light into the limited success MMP inhibitors have had in clinical settings as cancer therapeutics. Additionally, translating these insights into strategies to inhibit CAF-mediated matrix remodeling and subsequent CAF migration could offer novel avenues for metastasis prevention.

## 6. Acknowledgements

P.T.D., and P.E.B. gratefully acknowledge funding from the Delft Health Technology grant (between LUMC and TU Delft) and ZonMW grant (09120012010061). A.D.B. gratefully acknowledges funding from MSCA Postdoctoral Fellowships 2022 Project ID: 101111247.

## Notes

### Competing Interest Statement

The authors have declared no competing interest.

## References

[1] L. Wilkinson, T. Gathani, Understanding breast cancer as a global health concern, The British Journal of Radiology 95 (2021) 20211033–20211033. 10.1259/BJR.20211033.

[2] F. Lüönd, S. Tiede, G. Christofori, Breast cancer as an example of tumour heterogeneity and tumour cell plasticity during malignant progression, British Journal of Cancer 125 (2021) 164–175. 10.1038/s41416-021-01328-7.

[3] A. Muchlińska, A. Nagel, M. Popęda, J. Szade, M. Niemira, J. Zieliński, J. Skokowski, N. Bednarz-Knoll, A.J. Żaczek, Alpha-smooth muscle actin-positive cancer-associated fibroblasts secreting osteopontin promote growth of luminal breast cancer, Cellular and Molecular Biology Letters 27 (2022) 1–14. 10.1186/S11658-022-00351-7/FIGURES/3.

[4] G.W. van Pelt, T.P. Sandberg, H. Morreau, H. Gelderblom, J.H.J.M. van Krieken, R.A.E.M. Tollenaar, W.E. Mesker, The tumour–stroma ratio in colon cancer: the biological role and its prognostic impact, Histopathology 73 (2018) 197–206. 10.1111/HIS.13489.

[5] M. Xu, T. Zhang, R. Xia, Y. Wei, X. Wei, Targeting the tumor stroma for cancer therapy, Molecular Cancer 2022 21:1 21 (2022) 1–38. 10.1186/S12943-022-01670-1.

[6] D. Yan, X. Ju, B. Luo, F. Guan, H. He, H. Yan, J. Yuan, Tumour stroma ratio is a potential predictor for 5-year disease-free survival in breast cancer, BMC Cancer 22 (2022) 1–11. 10.1186/S12885-022-10183-5/FIGURES/4.

[7] N.M. Atallah, N. Wahab, M.S. Toss, S. Makhlouf, A.Y. Ibrahim, A.G. Lashen, S. Ghannam, N.P. Mongan, M. Jahanifar, S. Graham, M. Bilal, A. Bhalerao, S.E. Ahmed Raza, D. Snead, F. Minhas, N. Rajpoot, E. Rakha, Deciphering the Morphology of Tumor-Stromal Features in Invasive Breast Cancer Using Artificial Intelligence, Modern Pathology 36 (2023) 100254–100254. 10.1016/J.MODPAT.2023.100254.

[8] J. Wu, C. Liang, M. Chen, W. Su, Association between tumor-stroma ratio and prognosis in solid tumor patients: a systematic review and meta-analysis, Oncotarget 7 (2016) 68954–68954. 10.18632/ONCOTARGET.12135.

[9] R.M. Bremnes, T. Dønnem, S. Al-Saad, K. Al-Shibli, S. Andersen, R. Sirera, C. Camps, I. Marinez, L.T. Busund, The Role of Tumor Stroma in Cancer Progression and Prognosis: Emphasis on Carcinoma-Associated Fibroblasts and Non-small Cell Lung Cancer, Journal of Thoracic Oncology 6 (2011) 209–217. 10.1097/JTO.0B013E3181F8A1BD.

[10] N. Pasha, N.C. Turner, Understanding and overcoming tumor heterogeneity in metastatic breast cancer treatment, Nature Cancer 2 (2021). 10.1038/s43018-021-00229-1.

[11] B. Bareham, M. Dibble, M. Parsons, Defining and modeling dynamic spatial heterogeneity within tumor microenvironments, Current Opinion in Cell Biology 90 (2024) 102422–102422. 10.1016/J.CEB.2024.102422.

[12] G. Friedman, O. Levi-Galibov, E. David, C. Bornstein, A. Giladi, M. Dadiani, A. Mayo, C. Halperin, M. Pevsner-Fischer, H. Lavon, S. Mayer, R. Nevo, Y. Stein, N. Balint-Lahat, I. Barshack, H.R. Ali, C. Caldas, E. Nili-Gal-Yam, U. Alon, I. Amit, R. Scherz-Shouval, Cancer-associated fibroblast compositions change with breast cancer progression linking the ratio of S100A4+ and PDPN+ CAFs to clinical outcome, Nature Cancer 1 (2020) 692–708. 10.1038/s43018-020-0082-y.

[13] S.E. Reid, J. Pantaleo, P. Bolivar, M. Bocci, J. Sjölund, M. Morsing, E. Cordero, S. Larsson, M. Malmberg, B. Seashore-Ludlow, K. Pietras, Cancer-associated fibroblasts rewire the estrogen receptor response in luminal breast cancer, enabling estrogen independence, Oncogene 43 (2024) 1113–1126. 10.1038/S41388-024-02973-X.

[14] K.F. Goliwas, S. Libring, E. Berestesky, S. Gholizadeh, S.C. Schwager, A.R. Frost, T.R. Gaborski, J. Zhang, C.A. Reinhart-King, Mitochondrial transfer from cancer-associated fibroblasts increases migration in aggressive breast cancer, Journal of Cell Science 136 (2023). 10.1242/JCS.260419/VIDEO-3.

[15] N.I. Nissen, M. Karsdal, N. Willumsen, Collagens and Cancer associated fibroblasts in the reactive stroma and its relation to Cancer biology, Journal of Experimental and Clinical Cancer Research 38 (2019) 1–12. 10.1186/S13046-019-1110-6/TABLES/1.

[16] H. Li, Z. Qiu, F. Li, C. Wang, The relationship between MMP-2 and MMP-9 expression levels with breast cancer incidence and prognosis, Oncology Letters 14 (2017) 5865–5870. 10.3892/OL.2017.6924/HTML.

[17] J. Barbazán, D. Matic Vignjevic, Cancer associated fibroblasts: is the force the path to the dark side?, Current Opinion in Cell Biology 56 (2019) 71–79. 10.1016/J.CEB.2018.09.002.

[18] D. Pankova, Y. Chen, M. Terajima, M.J. Schliekelman, B.N. Baird, M. Fahrenholtz, L. Sun, B.J. Gill, T.J. Vadakkan, M.P. Kim, Y.H. Ahn, J.D. Roybal, X. Liu, E.R.P. Cuentas, J. Rodriguez, I.I. Wistuba, C.J. Creighton, D.L. Gibbons, J.M. Hicks, M.E. Dickinson, J.L. West, K.J. Grande-Allen, S.M. Hanash, M. Yamauchi, J.M. Kurie, Cancer-associated Fibroblasts Induce a Collagen Cross-link Switch in Tumor Stroma, Molecular Cancer Research : MCR 14 (2015) 287–287. 10.1158/1541-7786.MCR-15-0307.

[19] B. Erdogan, D.J. Webb, Cancer-associated fibroblasts modulate growth factor signaling and extracellular matrix remodeling to regulate tumor metastasis, Biochemical Society Transactions 45 (2017) 229–236. 10.1042/BST20160387.

[20] L. Liu, H. Yu, H. Zhao, Z. Wu, Y. Long, J. Zhang, X. Yan, Z. You, L. Zhou, T. Xia, Y. Shi, B. Xiao, Y. Wang, C. Huang, Y. Du, Matrix-transmitted paratensile signaling enables myofibroblast-fibroblast cross talk in fibrosis expansion, Proceedings of the National Academy of Sciences of the United States of America 117 (2020) 10832–10838. 10.1073/PNAS.1910650117/SUPPL_FILE/PNAS.1910650117.SM08.MP4.

[21] H. Su, M. Karin, M. Karin, Collagen architecture and signaling orchestrate cancer development, Trends in Cancer 9 (2023) 764–773. 10.1016/j.trecan.2023.06.002.

[22] B.K. Patel, K. Pepin, K.R. Brandt, G.L. Mazza, B.A. Pockaj, J. Chen, Y. Zhou, D.W. Northfelt, K. Anderson, J.M. Kling, C.M. Vachon, K.R. Swanson, M. Nikkhah, R. Ehman, Association of breast cancer risk, density, and stiffness: global tissue stiffness on breast MR elastography (MRE), Breast Cancer Research and Treatment 194 (2022) 79–79. 10.1007/S10549-022-06607-2.

[23] F. Calvo, N. Ege, A. Grande-Garcia, S. Hooper, R.P. Jenkins, S.I. Chaudhry, K. Harrington, P. Williamson, E. Moeendarbary, G. Charras, E. Sahai, Mechanotransduction and YAP-dependent matrix remodelling is required for the generation and maintenance of cancer-associated fibroblasts, Nature Cell Biology 2013 15:6 15 (2013) 637–646. 10.1038/ncb2756.

[24] A.K. Simi, M.F. Pang, C.M. Nelson, Extracellular matrix stiffness exists in a feedback loop that drives tumor progression, Advances in Experimental Medicine and Biology 1092 (2018) 57–67. 10.1007/978-3-319-95294-9_4/FIGURES/2.

[25] K.M. Riching, B.L. Cox, M.R. Salick, C. Pehlke, A.S. Riching, S.M. Ponik, B.R. Bass, W.C. Crone, Y. Jiang, A.M. Weaver, K.W. Eliceiri, P.J. Keely, 3D collagen alignment limits protrusions to enhance breast cancer cell persistence, Biophysical Journal 107 (2015) 2546–2558. 10.1016/j.bpj.2014.10.035.

[26] P.P. Provenzano, D.R. Inman, K.W. Eliceiri, S.M. Trier, P.J. Keely, Contact guidance mediated three-dimensional cell migration is regulated by Rho/ROCK-dependent matrix reorganization, Biophysical Journal 95 (2008) 5374–5384. 10.1529/biophysj.108.133116.

[27] P.P. Provenzano, K.W. Eliceiri, J.M. Campbell, D.R. Inman, J.G. White, P.J. Keely, Collagen reorganization at the tumor-stromal interface facilitates local invasion, BMC Medicine 4 (2006) 1–15. 10.1186/1741-7015-4-38/FIGURES/7.

[28] S.V. Bayer, W.R. Grither, A. Brenot, P.Y. Hwang, C.E. Barcus, M. Ernst, P. Pence, C. Walter, A. Pathak, G.D. Longmore, DDR2 controls breast tumor stiffness and metastasis by regulating integrin mediated mechanotransduction in cafs, eLife 8 (2019). 10.7554/ELIFE.45508.

[29] A. Marusyk, D.P. Tabassum, M. Janiszewska, A.E. Place, A. Trinh, A.I. Rozhok, S. Pyne, J.L. Guerriero, S. Shu, M. Ekram, A. Ishkin, D.P. Cahill, Y. Nikolsky, T.A. Chan, M.F. Rimawi, S. Hilsenbeck, R. Schiff, K.C. Osborne, A. Letai, K. Polyak, Spatial Proximity to Fibroblasts Impacts Molecular Features and Therapeutic Sensitivity of Breast Cancer Cells Influencing Clinical Outcomes, Cancer Research 76 (2016) 6495–6506. 10.1158/0008-5472.CAN-16-1457.

[30] Y. Wu, Y. Shi, Z. Luo, X. Zhou, Y. Chen, X. Song, S. Liu, Spatial multi-omics analysis of tumor-stroma boundary cell features for predicting breast cancer progression and therapy response, Front Cell Dev Biol 13 (2025) 1570696. 10.3389/fcell.2025.1570696.

[31] J. Ren, M. Smid, J. Iaria, D.C.F. Salvatori, H. Van Dam, H.J. Zhu, J.W.M. Martens, P. Ten Dijke, Cancer-associated fibroblast-derived Gremlin 1 promotes breast cancer progression, Breast Cancer Research : BCR 21 (2019) 109–109. 10.1186/S13058-019-1194-0.

[32] H.P.H. Naber, E. Wiercinska, P.T. Dijke, T. van Laar, Spheroid Assay to Measure TGF-β-induced Invasion, Journal of Visualized Experiments : JoVE (2011) 3337–3337. 10.3791/3337.

[33] V. Padmanaban, E.M. Grasset, N.M. Neumann, A.K. Fraser, E. Henriet, W. Matsui, P.T. Tran, K.J. Cheung, D. Georgess, A.J. Ewald, Organotypic culture assays for murine and human primary and metastatic-site tumors, Nature Protocols 15 (2020) 2413–2413. 10.1038/S41596-020-0335-3.

[34] N. Otsu, A Threshold Selection Method from Gray-Level Histograms, IEEE Transactions on Systems, Man, and Cybernetics 9 (1979) 62–66. 10.1109/TSMC.1979.4310076.

[35] R. Rezakhaniha, A. Agianniotis, J.T.C. Schrauwen, A. Griffa, D. Sage, C.V.C. Bouten, F.N. van de Vosse, M. Unser, N. Stergiopulos, Experimental investigation of collagen waviness and orientation in the arterial adventitia using confocal laser scanning microscopy, Biomech Model Mechanobiol 11 (2012) 461–473. 10.1007/s10237-011-0325-z.

[36] D. Böhringer, A. Bauer, I. Moravec, L. Bischof, D. Kah, C. Mark, T.J. Grundy, E. Görlach, G.M. O’neill, S. Budday, P. Strissel, R. Strick, A. Malandrino, R. Gerum, M. Mak, M. Rausch, B. Fabry, Fiber alignment in 3D collagen networks as a biophysical marker for cell contractility, (n.d.). 10.1101/2023.06.28.546896.

[37] L.M. Jawerth, S. Münster, D.A. Vader, B. Fabry, D.A. Weitz, A blind spot in confocal reflection microscopy: The dependence of fiber brightness on fiber orientation in imaging biopolymer networks, Biophysical Journal 98 (2010) L1–L3. 10.1016/j.bpj.2009.09.065.

[38] B. Davies, P.D. Brown, N. East, M.J. Crimmin, F.R. Balkwill, A synthetic matrix metalloproteinase inhibitor decreases tumor burden and prolongs survival of mice bearing human ovarian carcinoma xenografts, Cancer Res 53 (1993) 2087–2091.

[39] A. van der Net, Z. Rahman, A.D. Bordoloi, I. Muntz, P. Ten Dijke, P.E. Boukany, G.H. Koenderink, EMT-related cell-matrix interactions are linked to states of cell unjamming in cancer spheroid invasion, iScience 27 (2024) 111424. 10.1016/j.isci.2024.111424.

[40] C.D. Paul, P. Mistriotis, K. Konstantopoulos, Cancer cell motility: Lessons from migration in confined spaces, Nature Reviews Cancer 17 (2017) 131–140. 10.1038/nrc.2016.123.

[41] B. Trappmann, B.M. Baker, W.J. Polacheck, C.K. Choi, J.A. Burdick, C.S. Chen, Matrix degradability controls multicellularity of 3D cell migration, Nature Communications 8 (2017) 1–10.1038/s41467-017-00418-6.

[42] O. Ilina, P. Friedl, Mechanisms of collective cell migration at a glance, Journal of Cell Science 122 (2009) 3203–3208. 10.1242/jcs.036525.

[43] Z. Rahman, A.D. Bordoloi, H. Rouhana, M. Tavasso, G. van der Zon, V. Garbin, P. ten Dijke, P.E. Boukany, Interstitial flow potentiates TGF-β/Smad-signaling activity in lung cancer spheroids in a 3D-microfluidic chip, Lab on a Chip 24 (2024) 422–433. 10.1039/D3LC00886J.

[44] O. Ilina, P.G. Gritsenko, S. Syga, J. Lippoldt, C.A.M. La Porta, O. Chepizhko, S. Grosser, M. Vullings, G.J. Bakker, J. Starruß, P. Bult, S. Zapperi, J.A. Käs, A. Deutsch, P. Friedl, Cell–cell adhesion and 3D matrix confinement determine jamming transitions in breast cancer invasion, Nature Cell Biology 22 (2020) 1103–1115. 10.1038/s41556-020-0552-6.

