## Supporting Figures for "Inter-Spheroid Proximity and Matrix Remodeling Determine CAF-Mediated Cancer Cell Invasion"

### Supporting Information

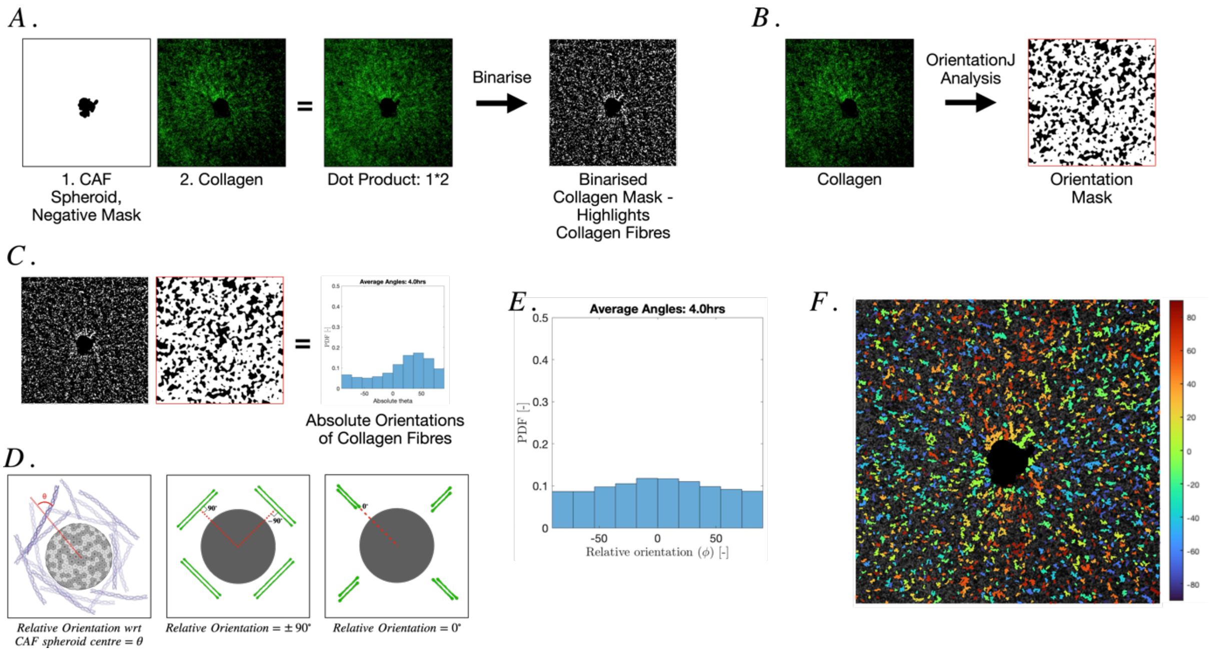


**Figure S1 – Analysis of caf mediated matrix remodelling**

A. Dot product of CAF spheroid negative mask and collagen fibre image to produce collagen mask with caf spheroid removed. This mask is further binarised to identify collagen fibers.
B. Collagen fibre image is run through OrientationJ plugin (vector field analysis, gaussian kernel size 10) to produce orientation mask.
C. Dot product of binarised collagen mask and Orientation mask is used to obtain absolute orientations of collagen fibers (with respect to x-axis).
D. Schematic of theta – relative orientation with respect to CAF spheroid centre. Based on OrientationJ and ImageJ conventions, collagen fibres oriented tangentially are ±90° and fibres oriented radially are 0°.
E. Histogram of relative collagen fibre orientation with its probability density function on the y-axis.
F. Representative image of caf spheroid and collagen channel overlaid with color-coded collagen fibers according to their relative orientation.

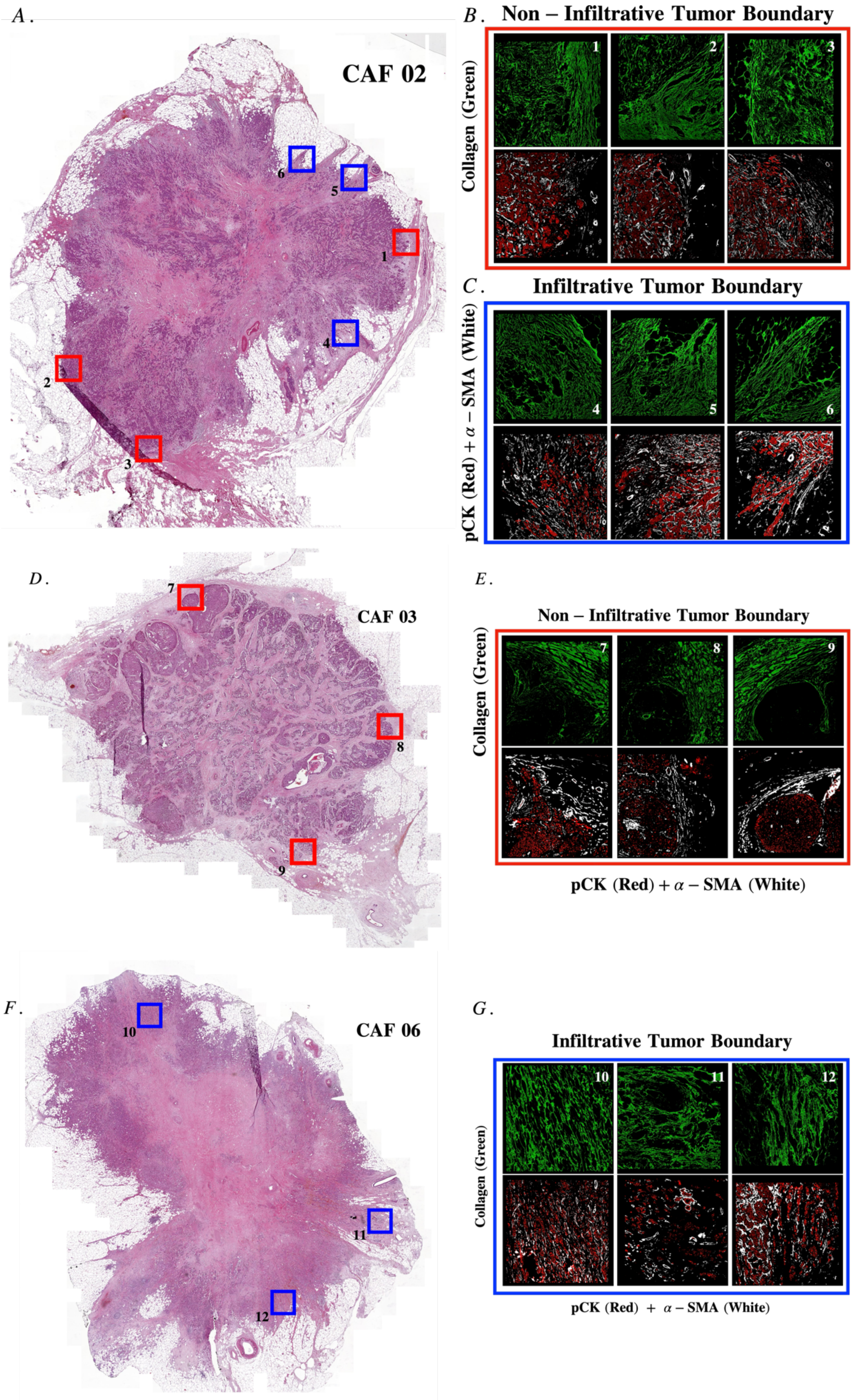
A.
CAF 02
1
2
3
4
5
6
B. Non – Infiltrative Tumor Boundary
Collagen (Green)
C. Infiltrative Tumor Boundary
pCK (Red) + α – SMA (White)
D.
CAF 03
7
8
9
E.
Non – Infiltrative Tumor Boundary
Collagen (Green)
pCK (Red) + α – SMA (White)
F.
CAF 06
10
11
12
G.
Infiltrative Tumor Boundary
Collagen (Green)
pCK (Red) + α – SMA (White)

**Figure S2 – Tumor tissue section of Tumor ID CAF02, CAF03 and CAF06 with highlighted Tumor boundaries and their respective zoomed in ROIs.**

A. Haematoxylin and Eosin (H&E) stained tissue section of Tumor ID CAF02 displaying both non-infiltrative (green) and infiltrative (blue) tumor boundaries.
B. Picrosirius red (PSR) stained tissue sections depicting collagen fibre bundles at non-infiltrative tumor boundaries. Multiplex immunofluorescence imaging of pCK (Red) and $\alpha\text{-}SMA$ (White) depicting cancer cell and CAF organization at non-infiltrative tumor boundaries.
C. PSR stained tissue sections depicting collagen fibre bundles at infiltrative tumor boundaries. Multiplex immunofluorescence imaging of pCK and $\alpha\text{-}SMA$ depicting cancer cell and CAF organization at infiltrative tumor boundaries.
D. H&E stained tissue section of Tumor ID CAF03 displaying only non-infiltrative tumor boundaries.
E. PSR stained tissue sections depicting collagen fibre bundles at non-infiltrative tumor boundaries. Multiplex immunofluorescence imaging of pCK (Red) and $\alpha\text{-}SMA$ (White) depicting cancer cell and CAF organization at non-infiltrative tumor boundaries.
F. H&E stained tissue section of Tumor ID CAF06 displaying only infiltrative tumor boundaries.
G. PSR stained tissue sections depicting collagen fibre bundles at infiltrative tumor boundaries. Multiplex immunofluorescence imaging of pCK and $\alpha\text{-}SMA$ depicting cancer cell and CAF organization at infiltrative tumor boundaries.

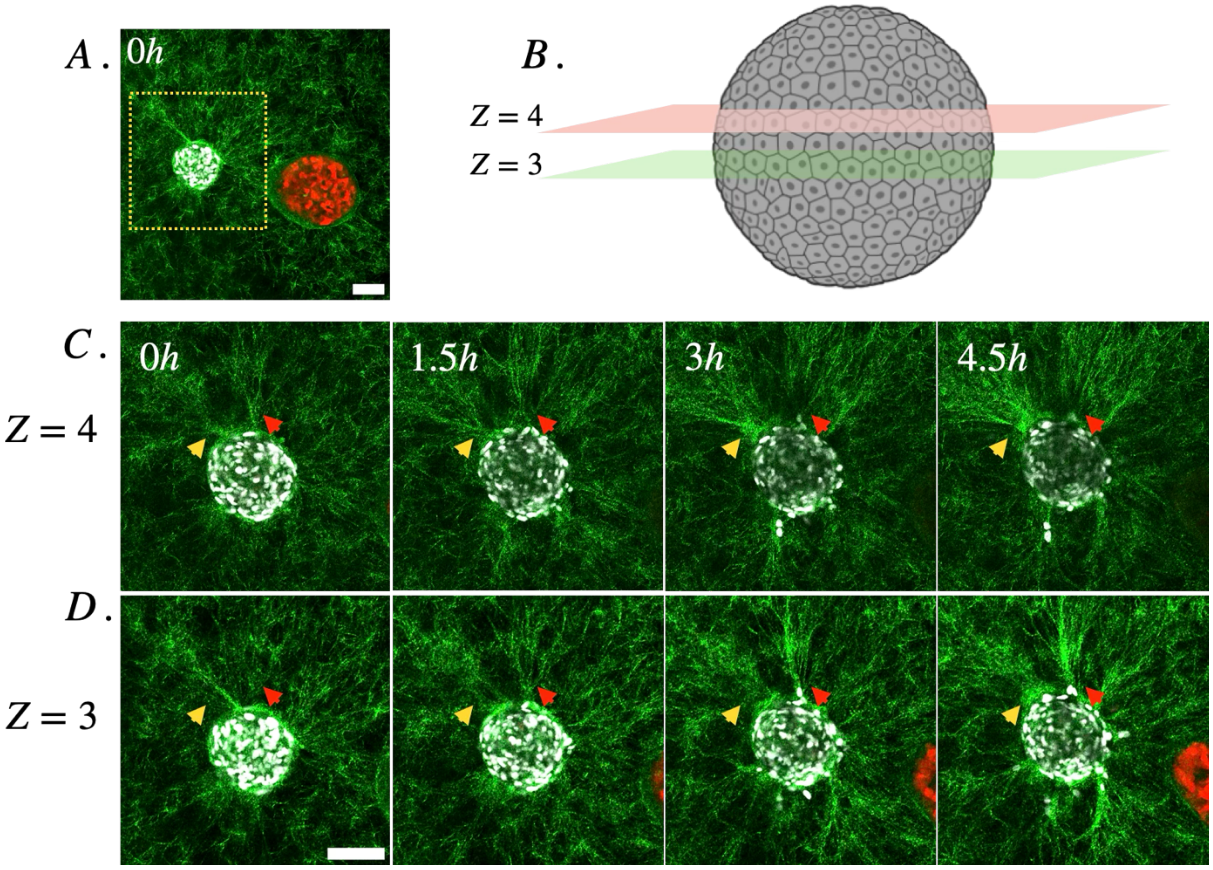


**Figure S3 – CAF spheriods aggregate collagen fibres from multiple focal planes to create thick collagen bundles**

A. Overview of 19TT-CAF and MCF7 cancer cell spheroids embedded in collagen matrix at T0. Yellow square highlights the ROI corresponding to C and D.
B. Visualization of consecutive Z focal planes 3 and 4. Each slice is 20μm thick.
*C.* CAF and Collagen visualized at *Z* = *4*. Yellow arrows correspond to the collagen fibres that are pulled into the imaging plane at *Z* = *4* while Red arrows show collagen fibres that are pulled out of the *Z* = *4* focal plane.
*D.* CAF and Collagen visualized at *Z* = *3*. Yellow arrows correspond to the collagen fibres that are pulled out of the focal plane at *Z* = *3* while Red arrows show collagen fibres that are pulled out of the *Z* = *3* imaging plane.

Scale Bar = 100 μm

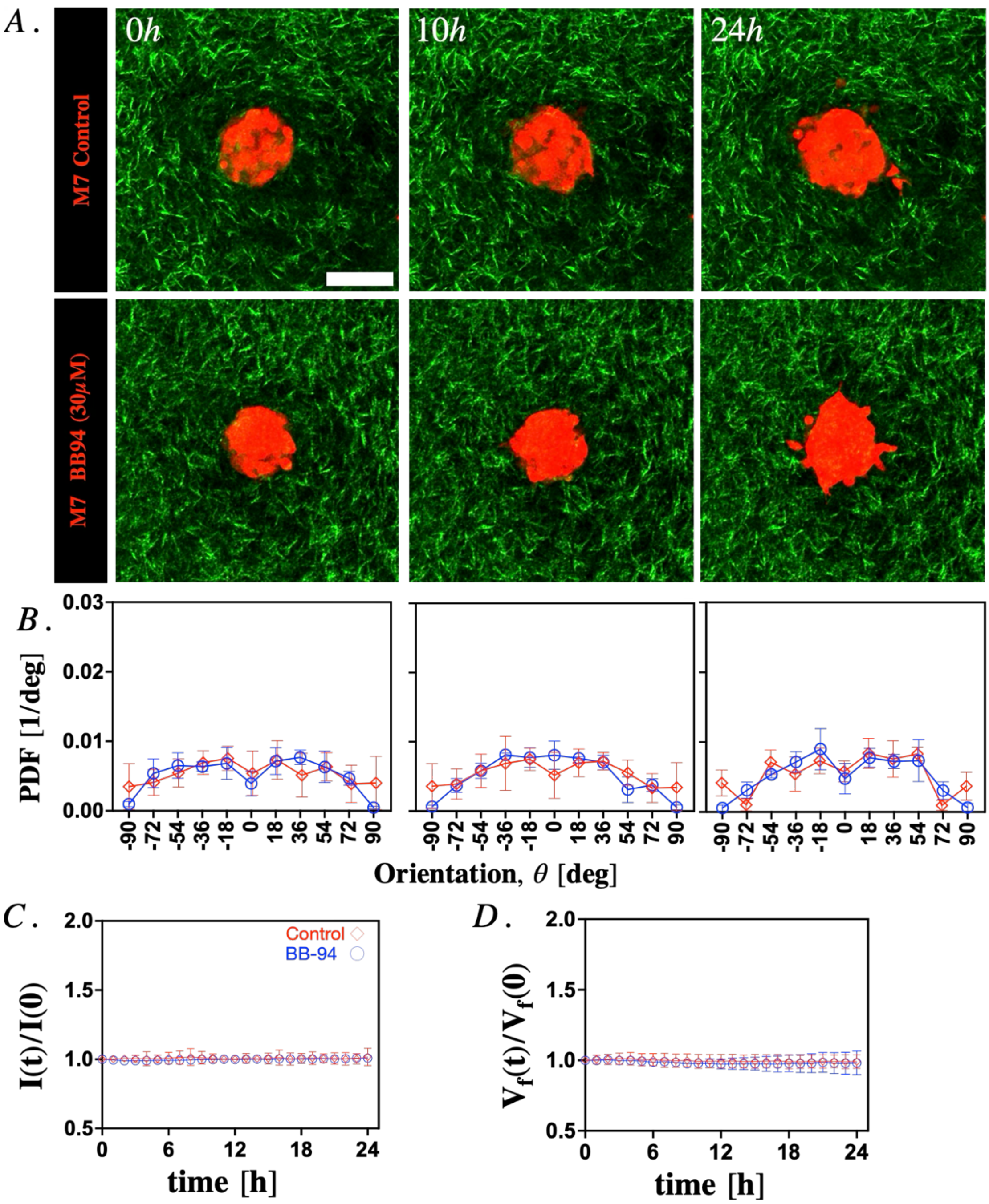


**Figure S4 – M7 spheroids embedded in collagen matrix treated with broad spectrum MMP inhibitors (BB94, 30 μM)**

A. Representative time-lapse images of Vehicle Control and BB94 treated MCF7 spheroids in collagen matrix at 0, 10 and 24 hours.
B. Relative orientation of collagen fibres with respect to MCF7 spheroid centroid at time 0, 10 and 24 hours. Veh Control (n = 12) and BB94 treated (n = 5) graphs are in Red and Blue respectively.
C. Pixel Intensity of collagen fibres over 24 hours. Veh Control (n = 12) and BB94 treated (n = 5) graphs are in Red and Blue respectively.
D. $V_f$ of collagen ECM over 24 hours. Veh Control (n = 12) and BB94 treated (n = 5) graphs are in Red and Blue respectively.

Scale Bar = 100 μm

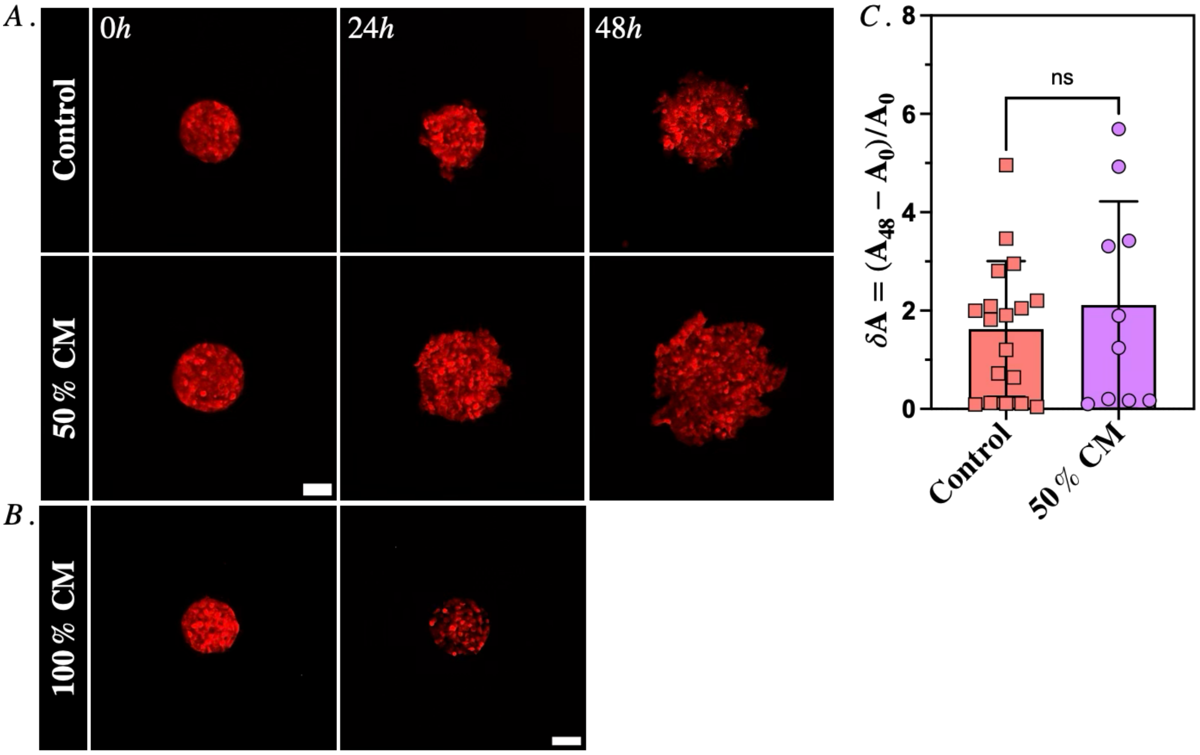


**Figure S5 - Stimulation by CAF Conditioned Medium (CM) does not have a statistically significant effect on M7 spheroid expansion.**

A. M7 spheroids embedded in collagen matrices were stimulated with CM obtained from CAF culture. As shown in the figure, 50% CM (1:1 dilution of conditioned medium in culture medium) had little effect on M7 spheroid expansion and no effect on M7 single cell dissemination compared to negative controls.
B. The use of 100% CM on M7 spheroids resulted in significant cell death after 24 h and thus were excluded from further analysis.
C. Quantification of M7 spheroid expansion ($n = 18$ for control, $n = 9$ for 50% CM) showed, on average, a slightly higher peak for M7 spheroids stimulated with CM but results were highly inconsistent with M7 spheroids control also showing similar behaviour ($p = 0.7116$).

Scale Bar = 100 μm

#### Movie Captions

Movie 1. 48 hr Time-lapse of MCF7 spheroid embedded in 3mg/ml collagen matrix.

Movie 2. 48 hr time-lapse of 19TT CAF and MCF7 spheroids embedded in 3mg/ml collagen matrix separated by a distance of $d \geq D\_C$.

Movie 3. 48 hr time-lapse of 19TT CAF and MCF7 spheroids embedded in 3mg/ml collagen matrix separated by a distance of $d < D\_C$.

Movie 4. 24 hr time-lapse video of 19TT CAF spheroid remodeling 3mg/ml collagen matrix.

Movie 5. 24 hr time-lapse video of MCF7 spheroid in collagen matrix.

Movie 6. 24 hr time-lapse video of 19TT CAF spheroid embedded in collagen matrix and treated with Batimastat (BB-94, 30μM).

Movie 7. 24 hr time-lapse video of Veh Control 19TT CAF spheroid embedded in collagen matrix.

Movie 8. 48 hr time-lapse of MCF7 and 19TT CAF spheroid embedded in collagen matrix treated with Batimastat (BB-94, 30μM).

Movie 9. 48 hr time-lapse of 19TT CAF and MCF7 heterospheorid embedded in collagen matrix and treated with Batimastat (BB-94, 30μM).

Movie 10. 48-hr time-lapse of Veh Control 19TT CAF and MCF7 heterospheorid embedded in collagen matrix.

Movie 11. 48-hr time-lapse of 19TT CAF and MCF7 heterospheorid embedded in collagen matrix treated with Batimastat (BB-94, 30μM).
